## Supplementary Figures for "Population-scale long-read sequencing uncovers transposable elements contributing to gene expression variation and associated with adaptive signatures in *Drosophila melanogaster*"

**Figure S1**. Scatter plots showing the correlation between: Top Row (yellow dots): Sequenced gigabases (Gb) and **A)** Genome Assembly size in Mb (GenomeSize), **B)** TE copies and **C)** TE content (%). Middle Row (blue dots): Read Length (N50) and **D)** Genome Assembly size in Mb (GenomeSize) and **E)** TE content (%). Bottom Row (green dots): Genome N50 (Mb) and **F)** Complete BUSCO (%), **G)** TE copies and **H)** TE content (%). $R^2$ values indicate the coefficient of determination for each pair of variables.

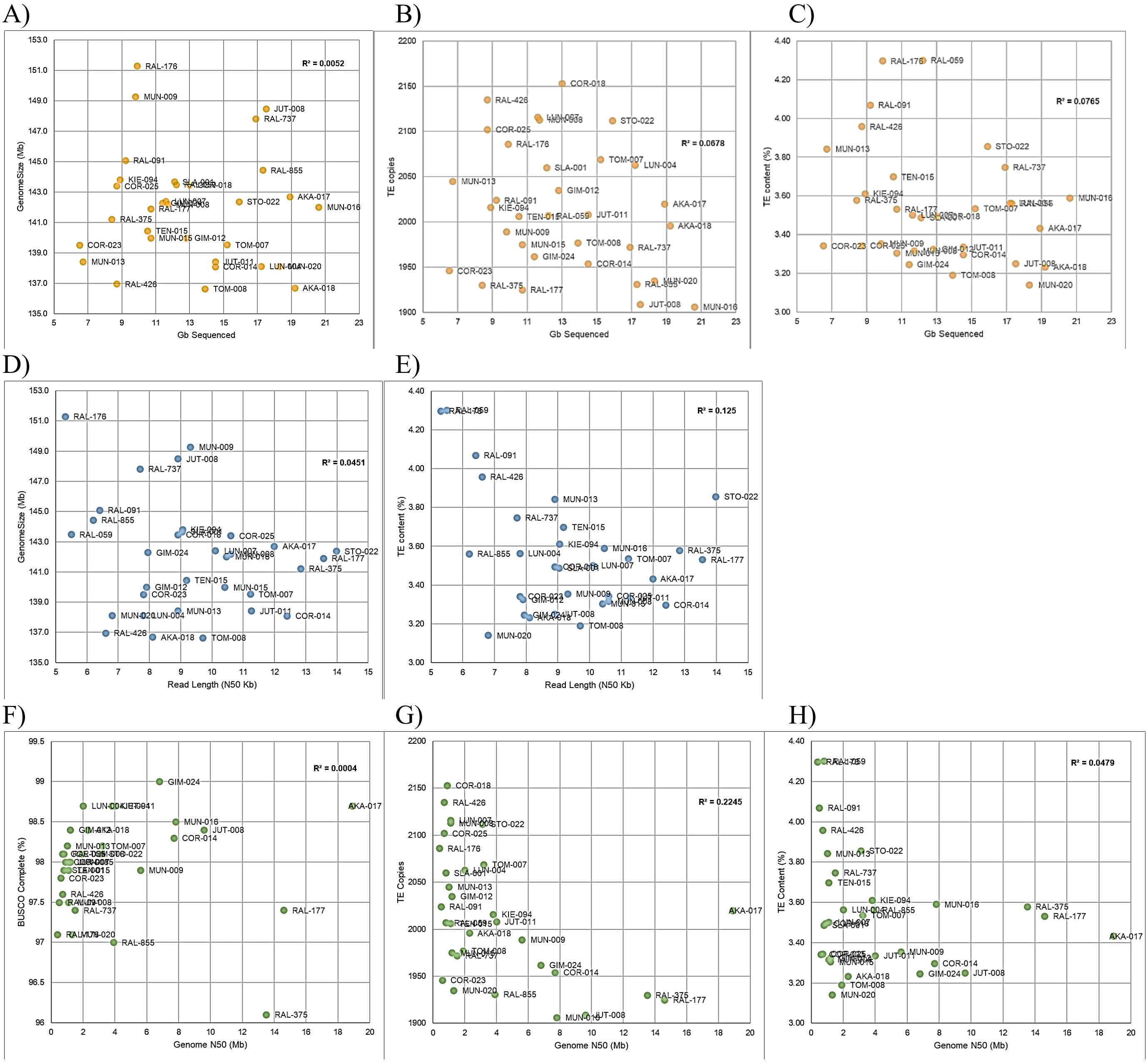

**Figure S2.** Scatter plot showing the relationship between the percentage of genomes in which the piRNA clusters were detected (y axis) and the size (in Kb) of the piRNA clusters (x axis). Pearson correlation=-0.47.

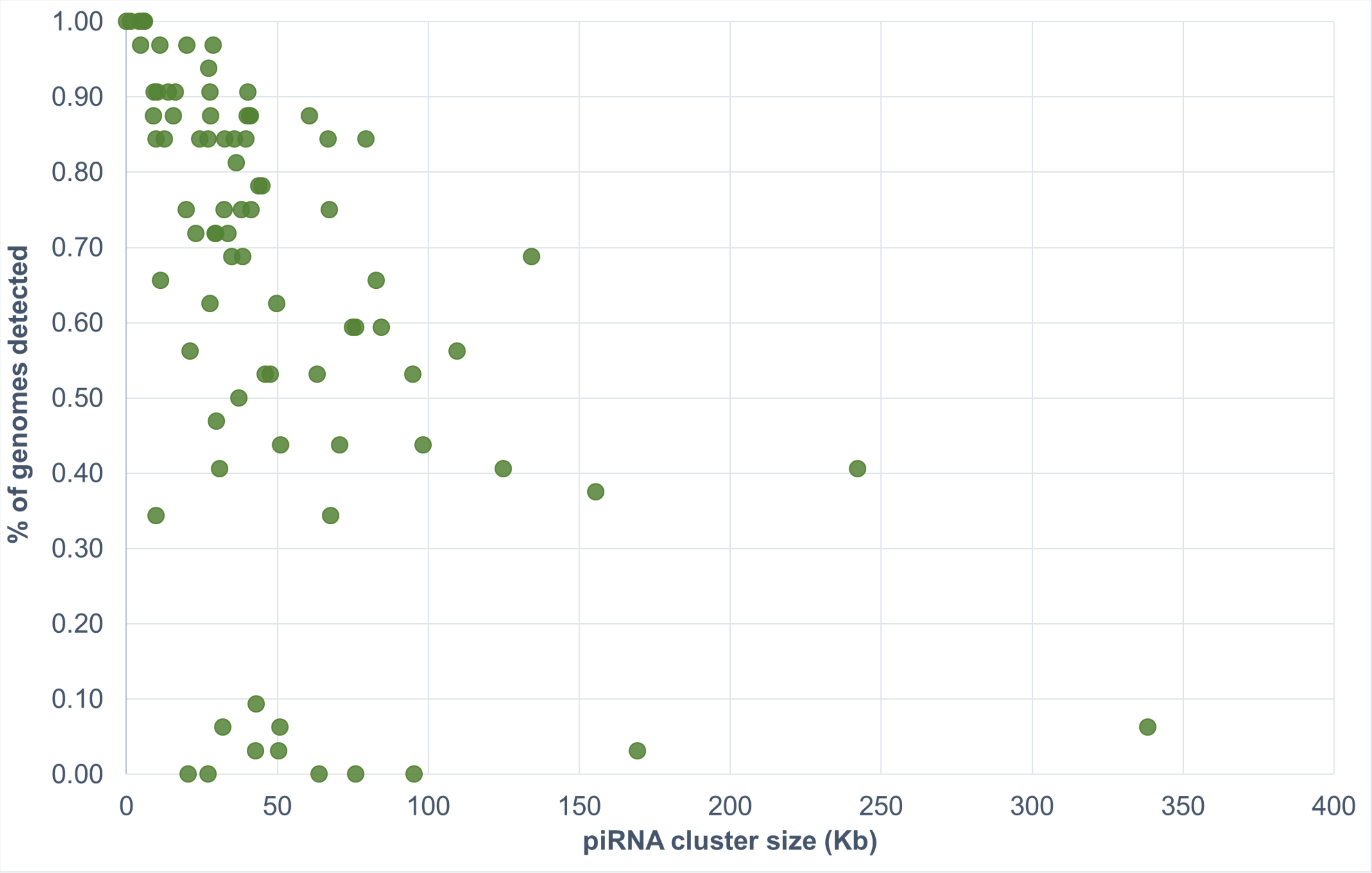

**Figure S3.** Dot plots showing the alignment of each sequenced genome (y axis) against the major chromosomal arms of the ISO1 genome (x axis).

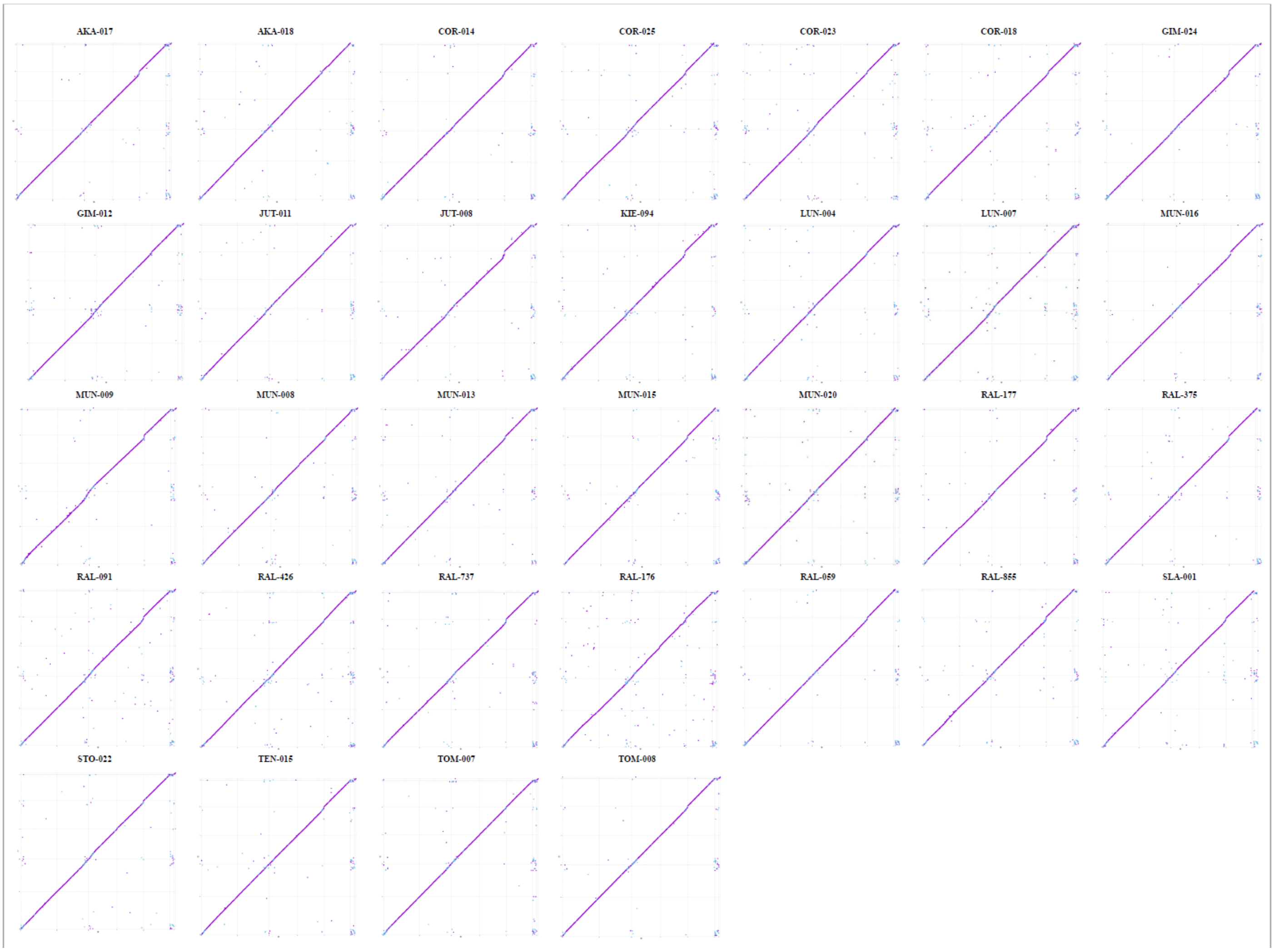

**Figure S4.** Sequence alignments of two newly identified TE families. **A)** Multiple Sequence Alignment of all **NewFam14 (MITE)** copies in three of the sequenced genomes. **B)** Pairwise alignment between consensus sequence of **NewFam06 (TIR)** and its closest sequence in RepBase (*EnSpm-1_JC*), a DNA transposon from the *Jatropha curcas* genome. The sequences share a 51.4% identity if only the aligned region is considered (local alignment).

A)

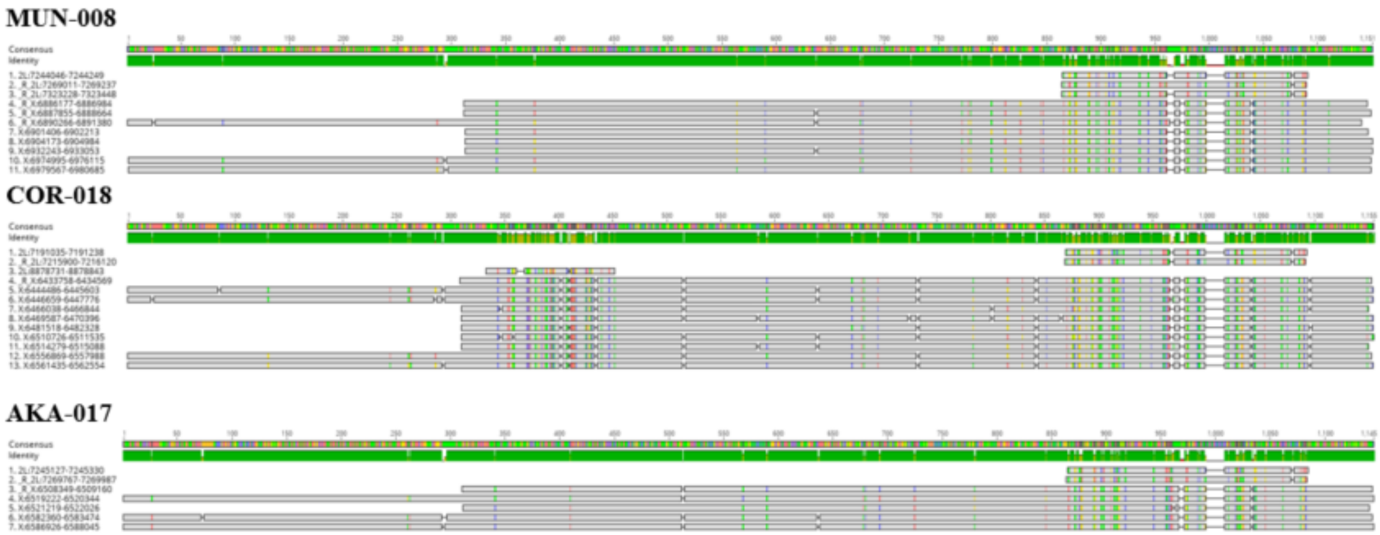


B)

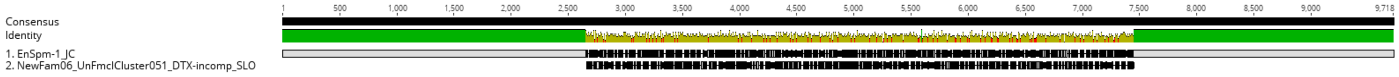

**Figure S5.** Percentage of sequence identity of the copies in the 32 genomes and in the ISO1 reference genome for each of the three new families.

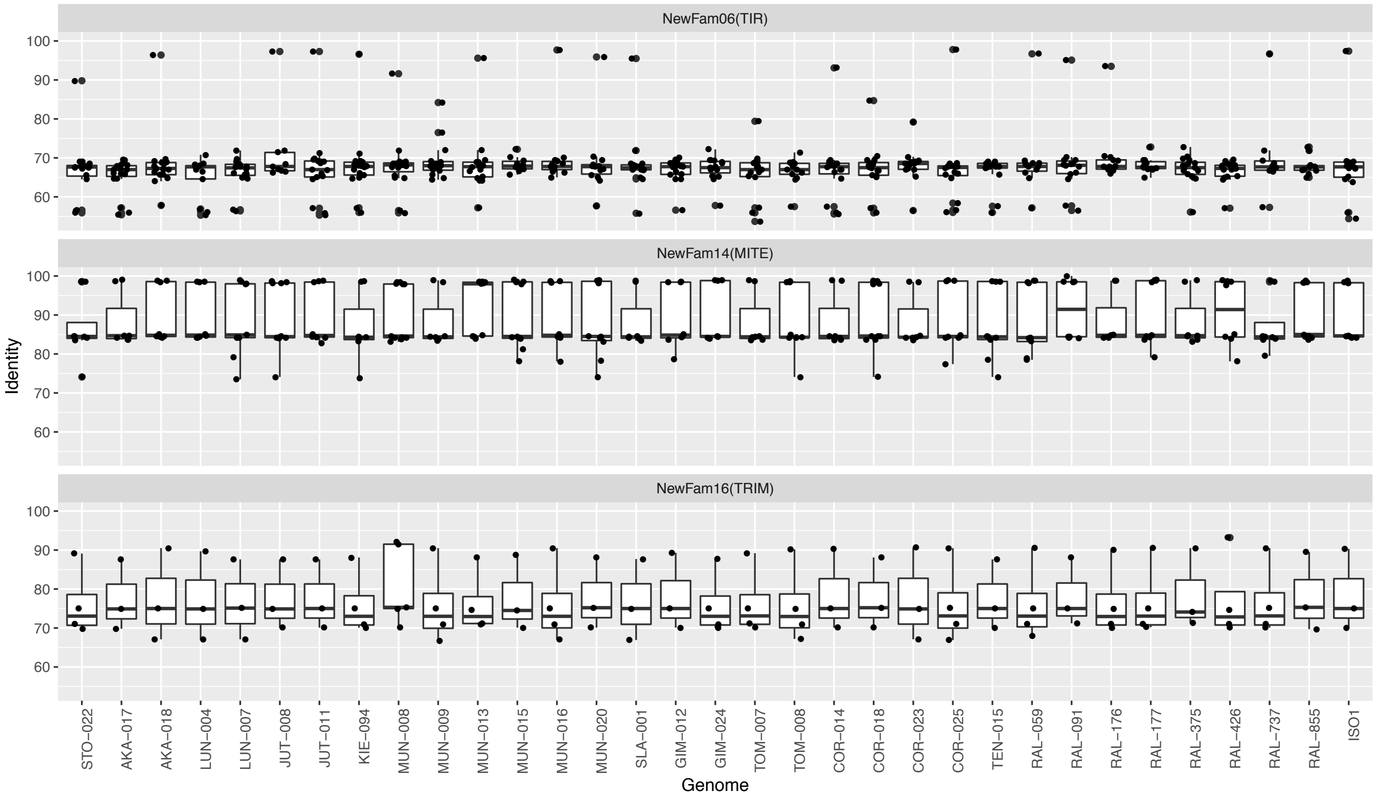

**Figure S6.** Proportion of TE Orders annotated in each genome. **A)** Base pairs occupied. **B)** Number of TE copies.

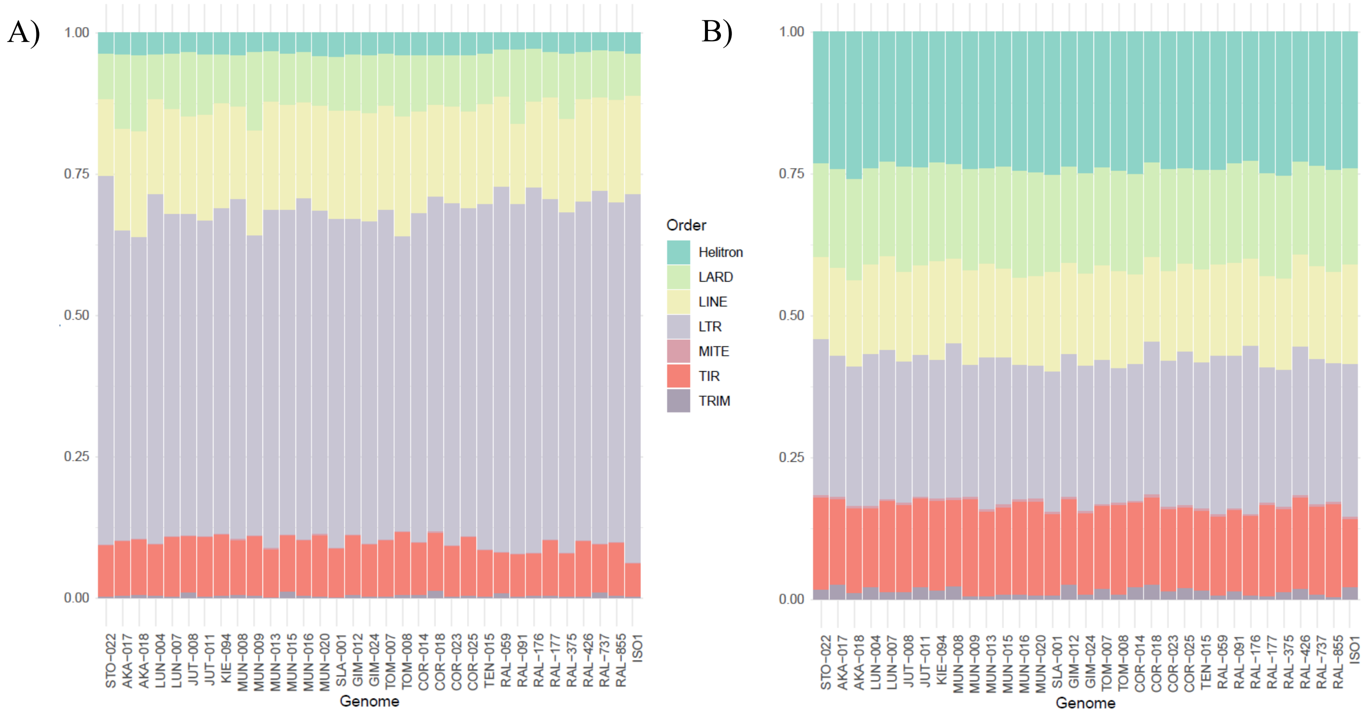

**Figure S7.** Clustering and heatmap representing the number of TE copies per family in the 32 genomes plus the ISO1 reference. Colors represent family Z-score. Clustering distance: correlation; clustering method: average.

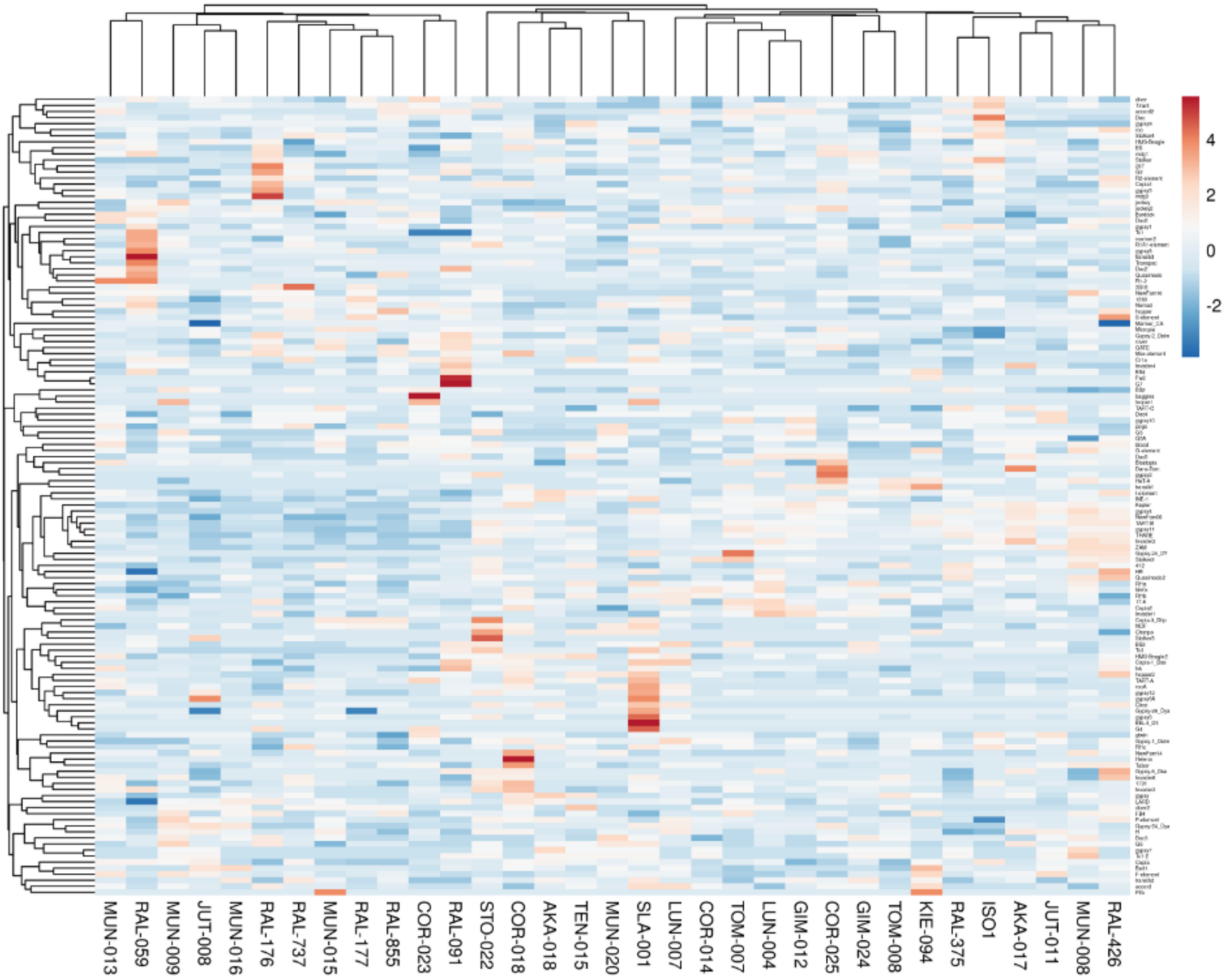

**Figure S8.** Heatmap representing the number of copies in the 20 most variable families with at least 10 copies in one strain. Colors represent family Z-score.

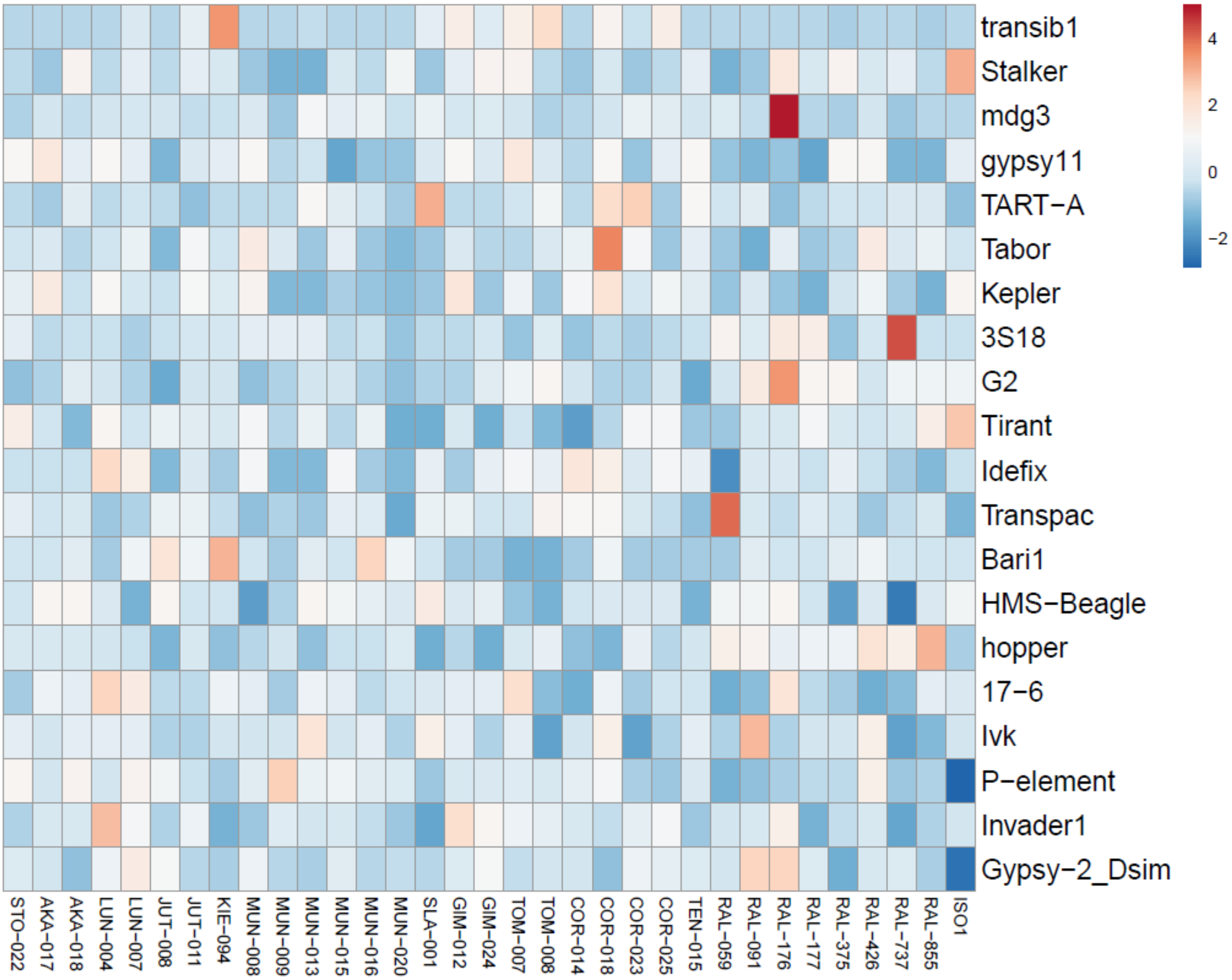

**Figure S9.** Boxplots showing the distribution of sequence identity values for copies from different superfamilies in each genome.

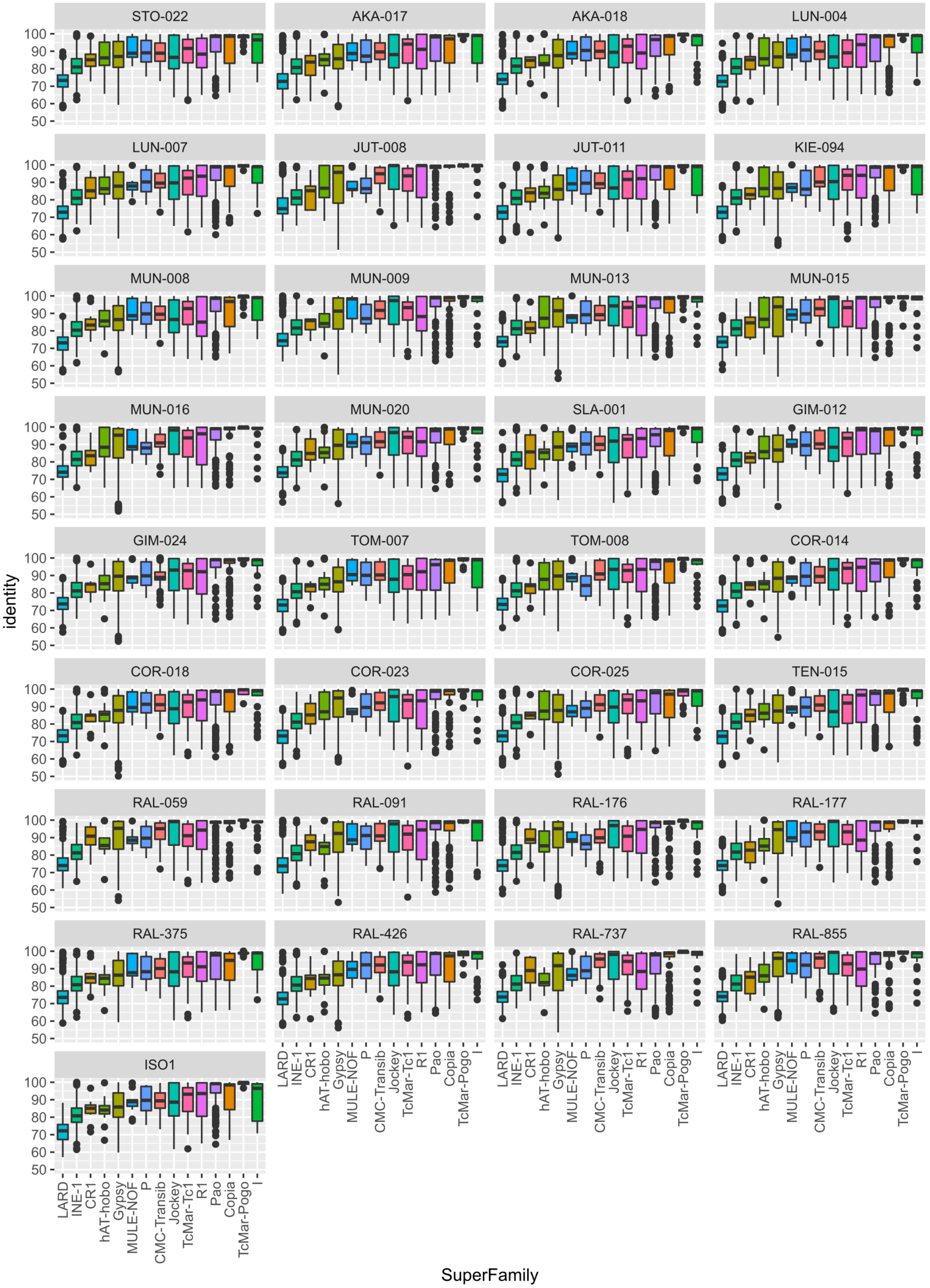

**Figure S10.** Distribution of TE identity values**. A).** Pairwise comparison in the distribution of TE identity values. Colors in the matrix represent adjusted p-values (Kolmogorov-Smirnov's test) when comparing the distributions of identities between genomes. **B)** Significance of pairwise Wilcoxon test for identity values in *Gypsy* and *Jockey* copies. Dot size and color-scale represent the –log10(adjusted p-value). Note adjusted p-value < 0.05 correspond with –log10(adjusted p-value) > 1.3.

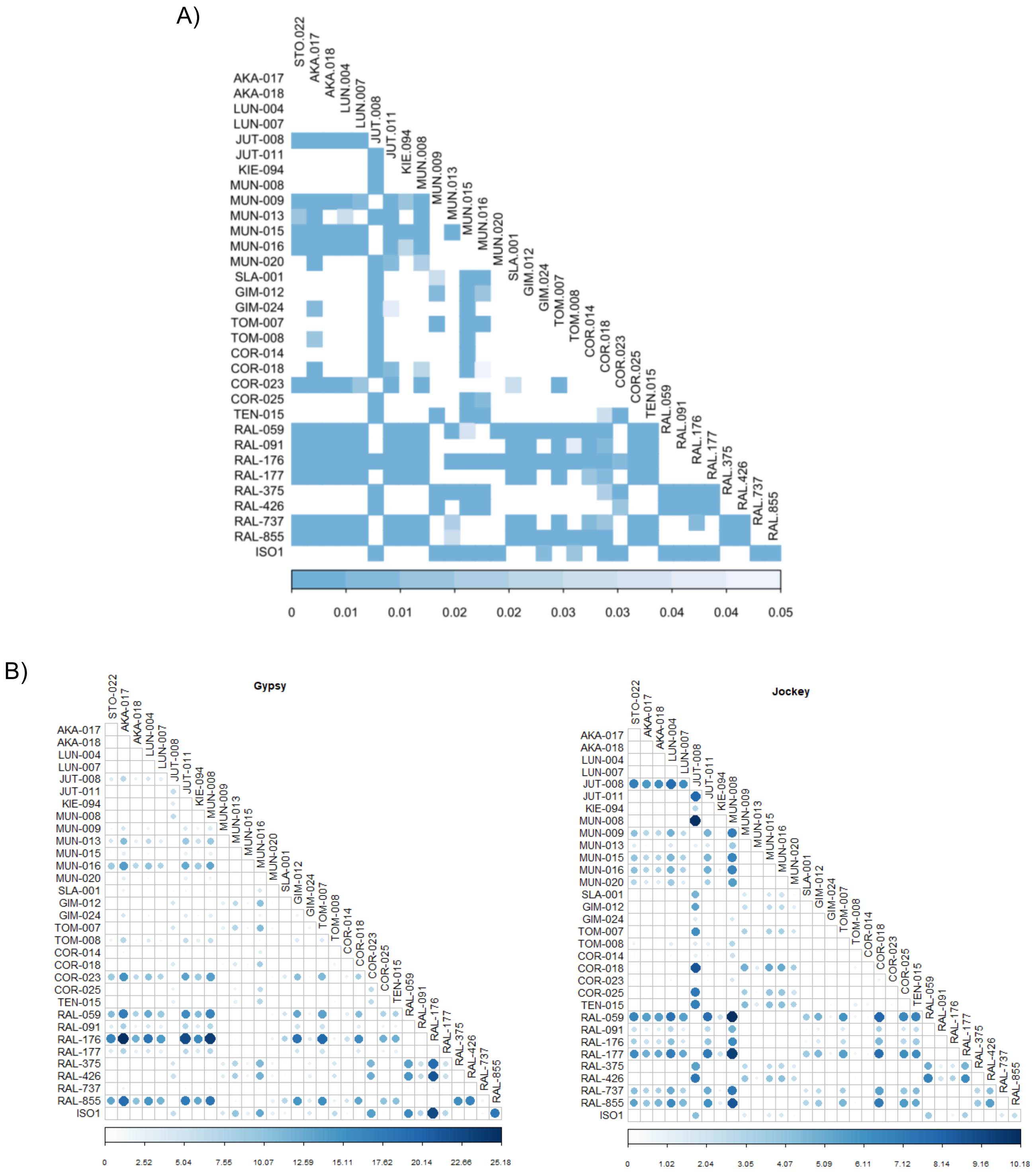

**Figure S11.** SFS for TE insertions and for SNPs with high, moderate, and low predicted effect according to SnpEff tool and for SNPs located in introns and considered to be evolving neutrally**.** Pairwise Kolmogorov-Smirnov test for SFS distributions for different types of variants showed significant differences for all comparisons ($p<0.0001$). However, *D* statistics showed greater differences between TEs vs. LOW impact (D= 0.70544) and TEs vs. Neutral SNPs (D=0.70415) than TEs vs, HIGH/MODERATE impact SNPs (D= 0.49492 and 0.59761, respectively), which is also consistent with a deleterious or slightly deleterious effect of TEs.

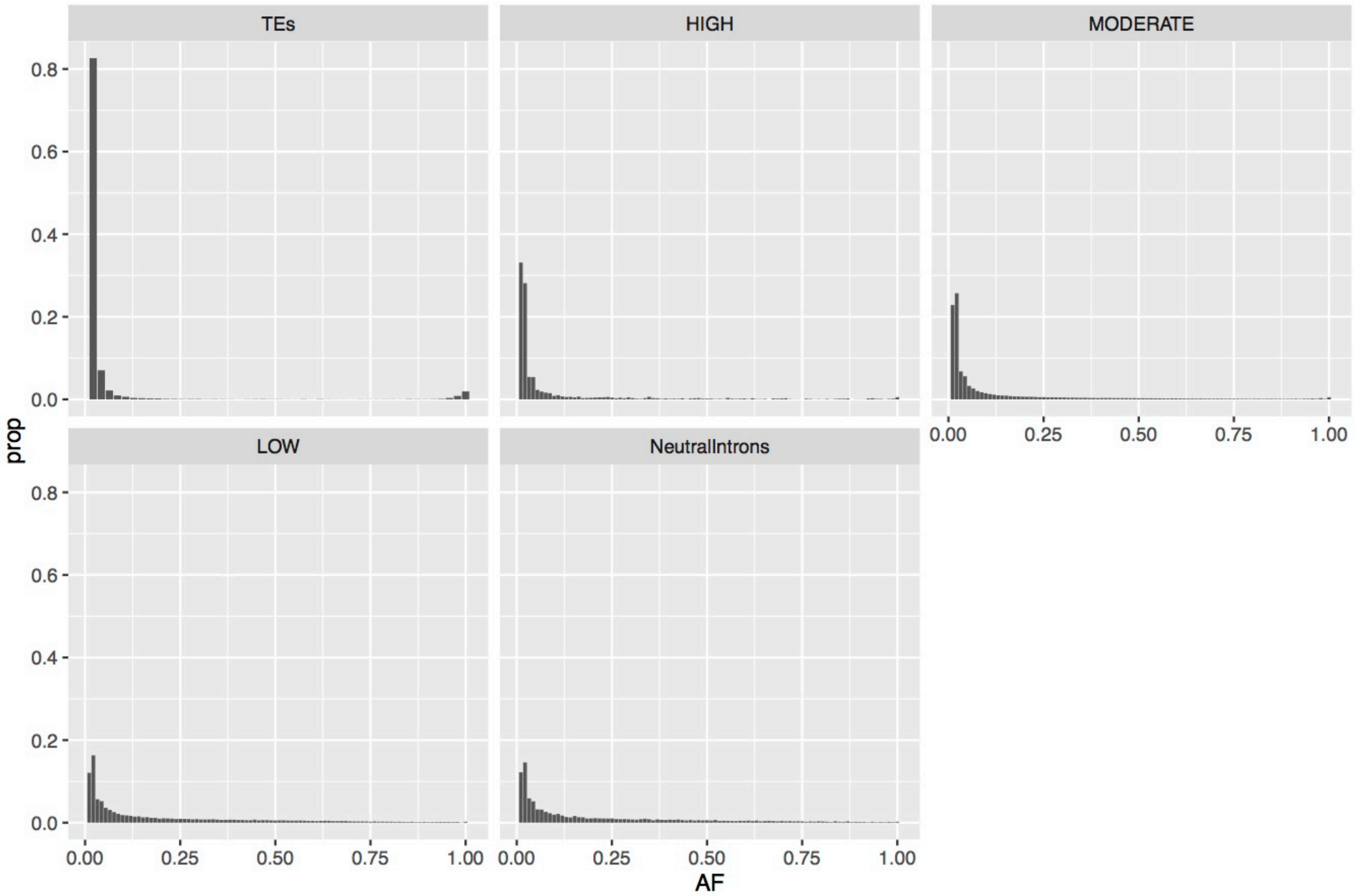

**Figure S12.** Schematic representation of the pipeline applied for the *de novo* genome assembly of the PacBio (**A**) and the Nanopore (**B**) sequences. Green boxes represent the software used, orange represent the input files and grey the resulting output from each step.

**A)**

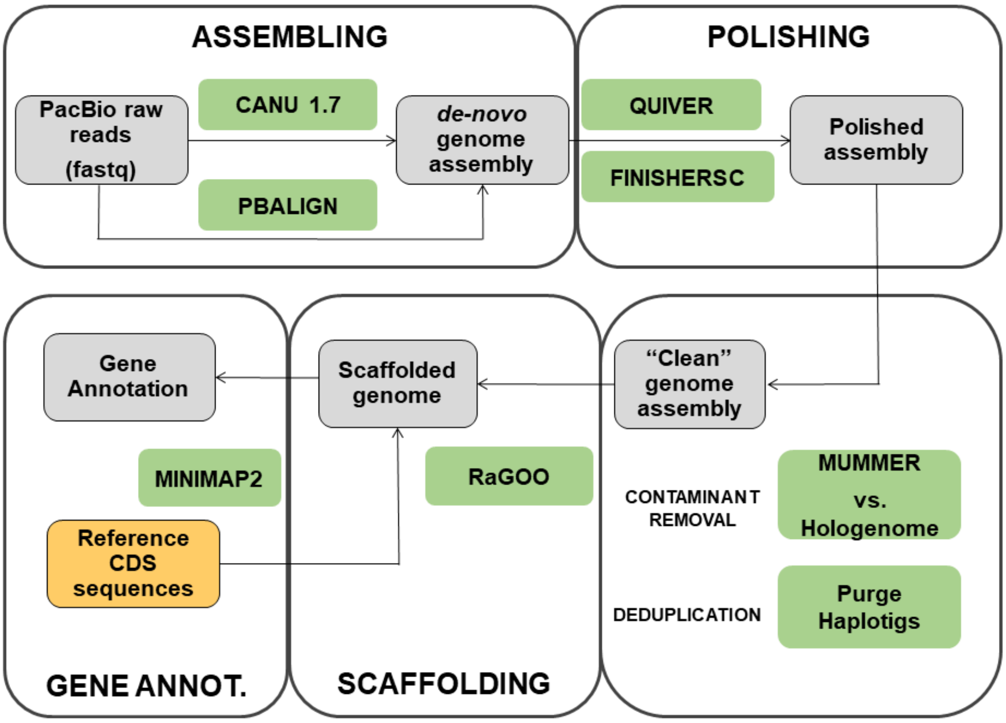


**B)**

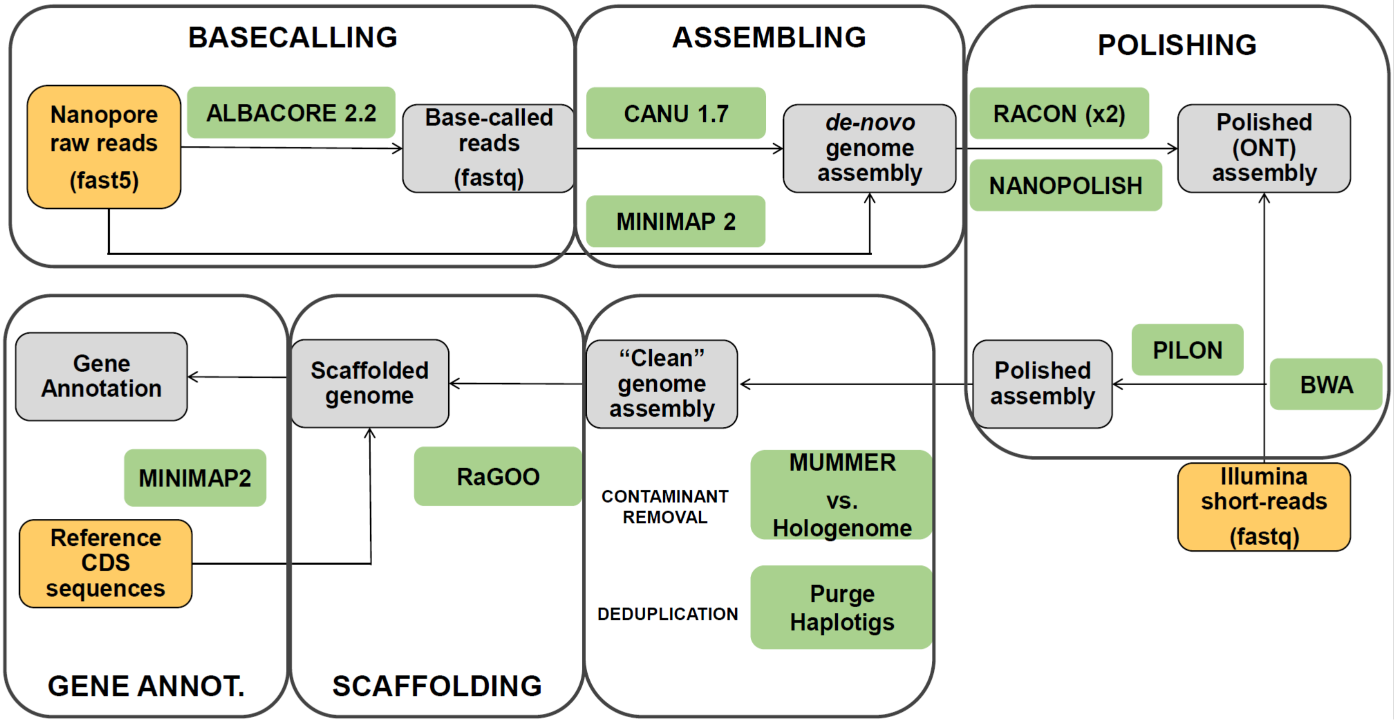

**Figure S13.** Scatter plot showing the relationship between the percentage of repeats in the genome according to *RepeatMasker* (% Repeats) and the size of the raw assemblies in Mb (SizeMb). The size of the dots represents the N50 (in Mb) of de contigs.

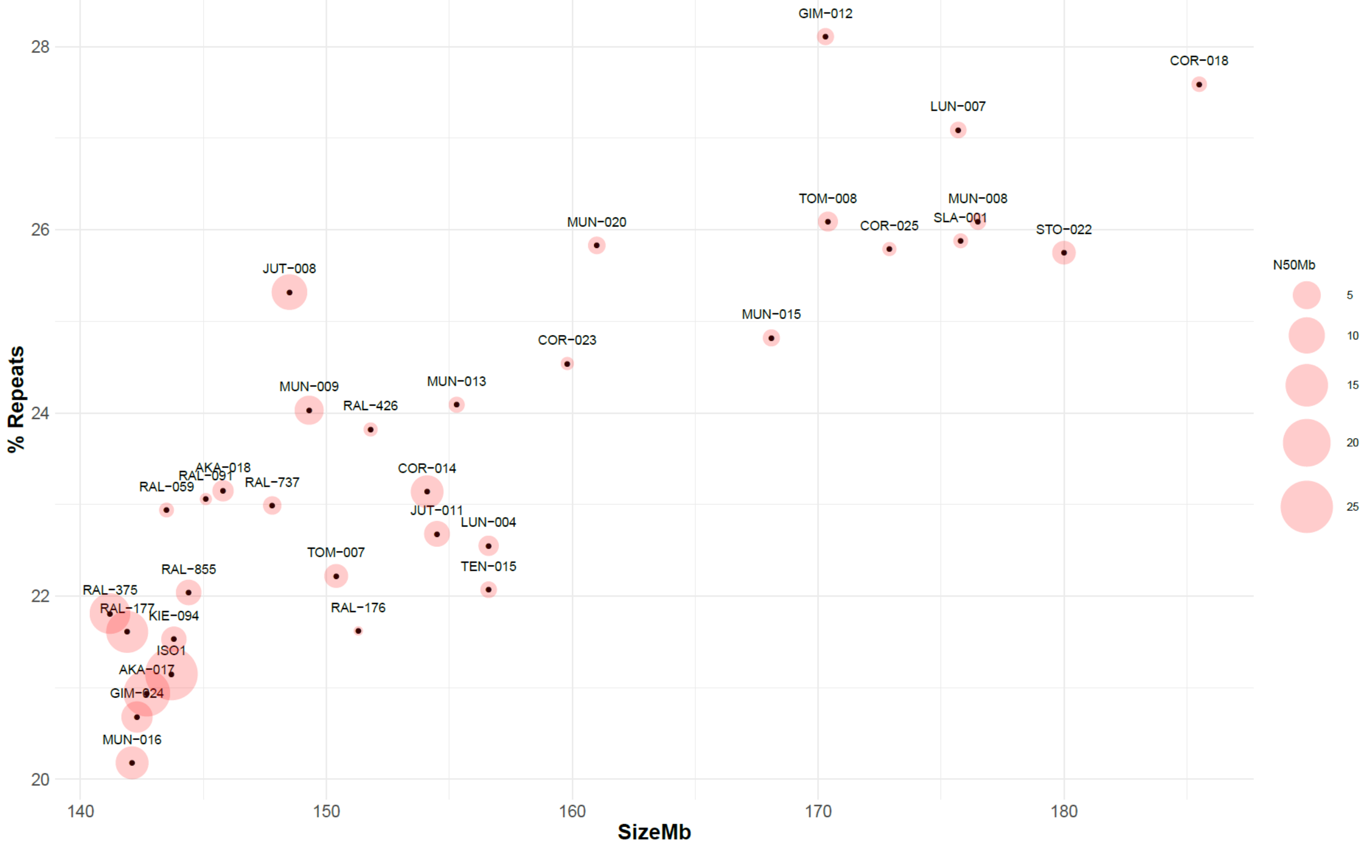

**Figure S14.** Scatter plot showing the relationship between percentage of repeats in the genome according to *RepeatMasker* (% Repeats,) and the genome assembly N50 (Genome N50, in Mb).

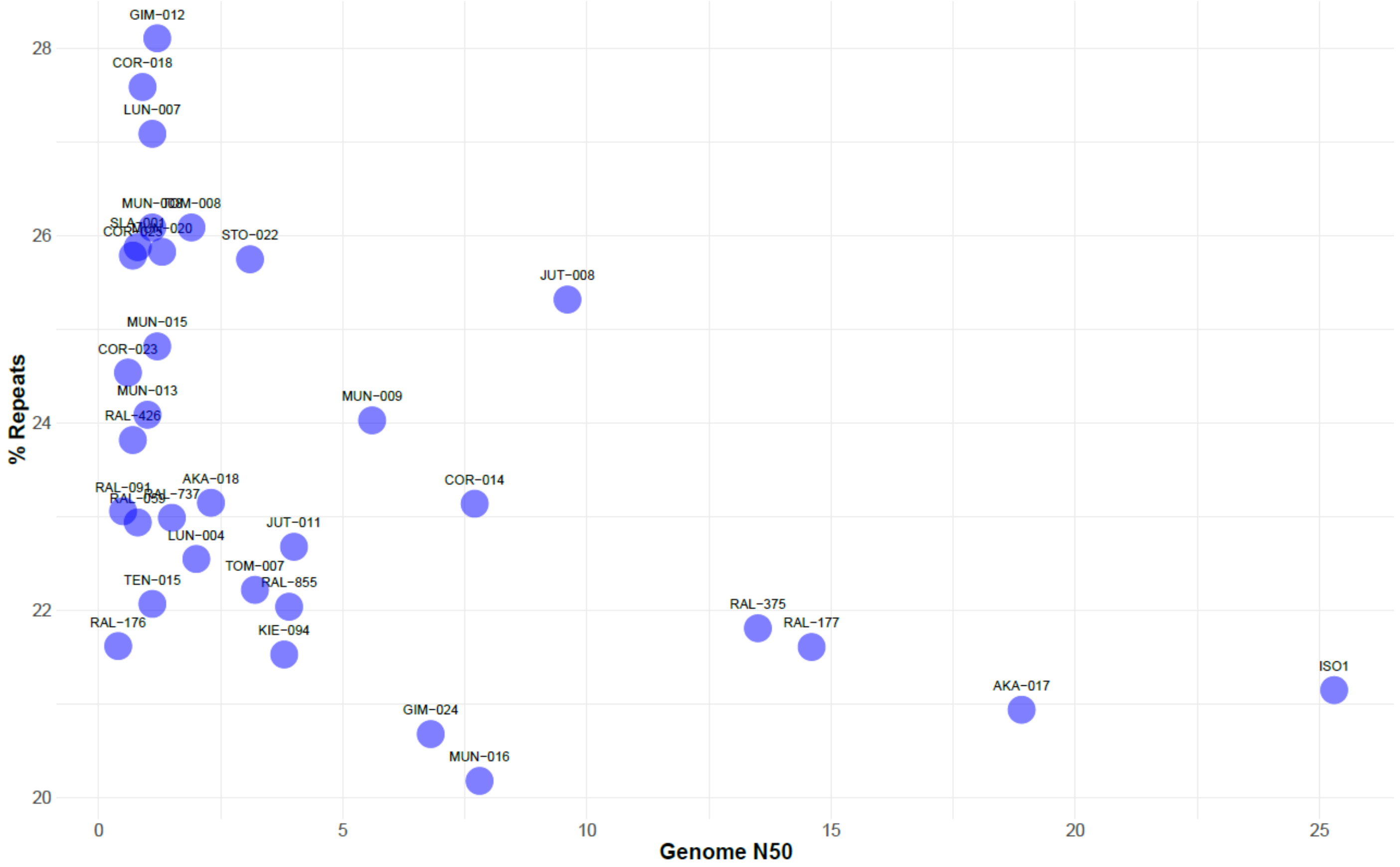

**Figure S15.** Scatter plot showing the relationship between the percentage of *BUSCO* duplicates and the size of the raw assemblies in Mb (SizeMb).

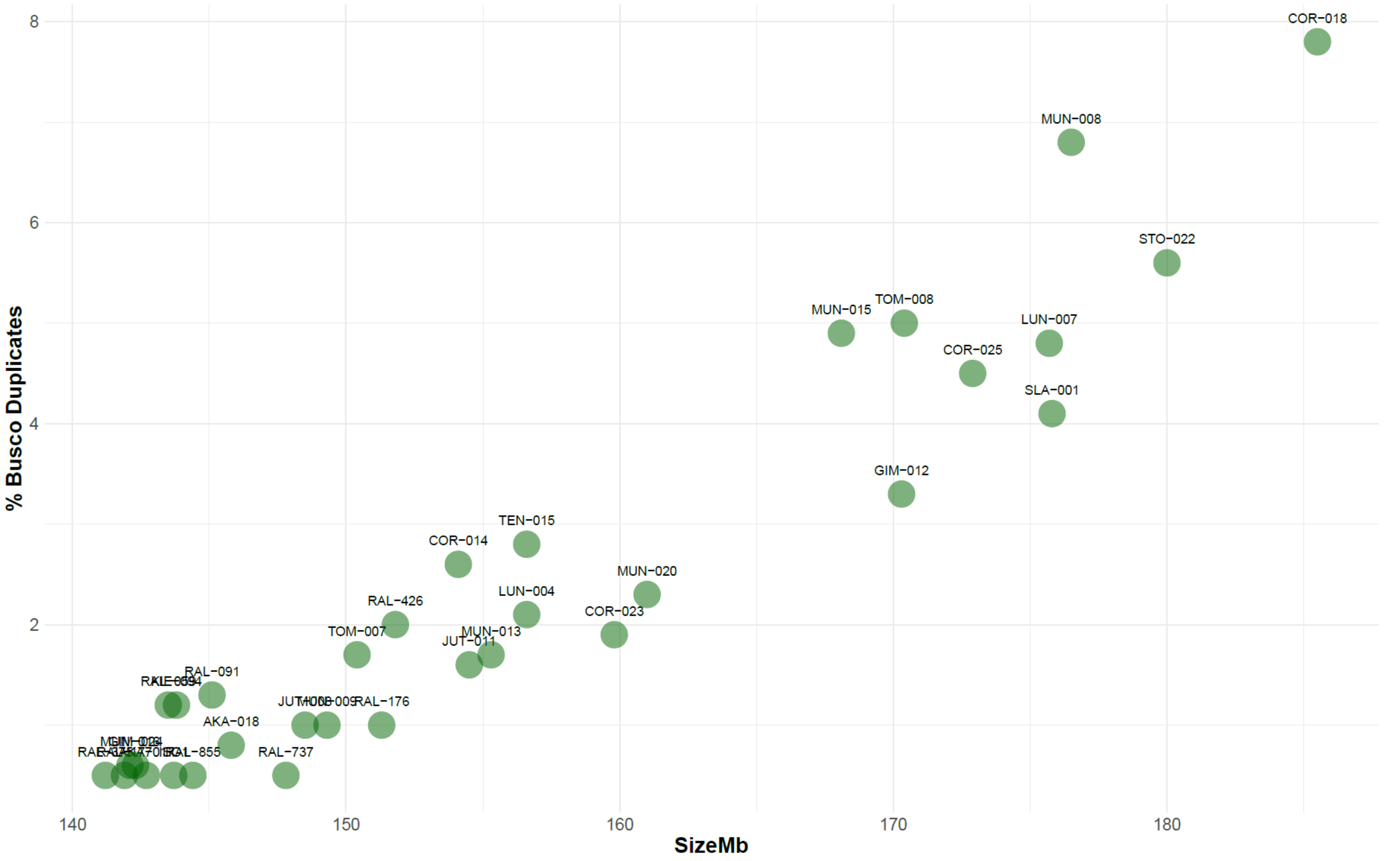

**Figure S16.** Scatter plot showing the relationship between the percentage of heterozygosity calculated using Illumina sequences and the size of the raw assemblies in Mb (SizeMb).

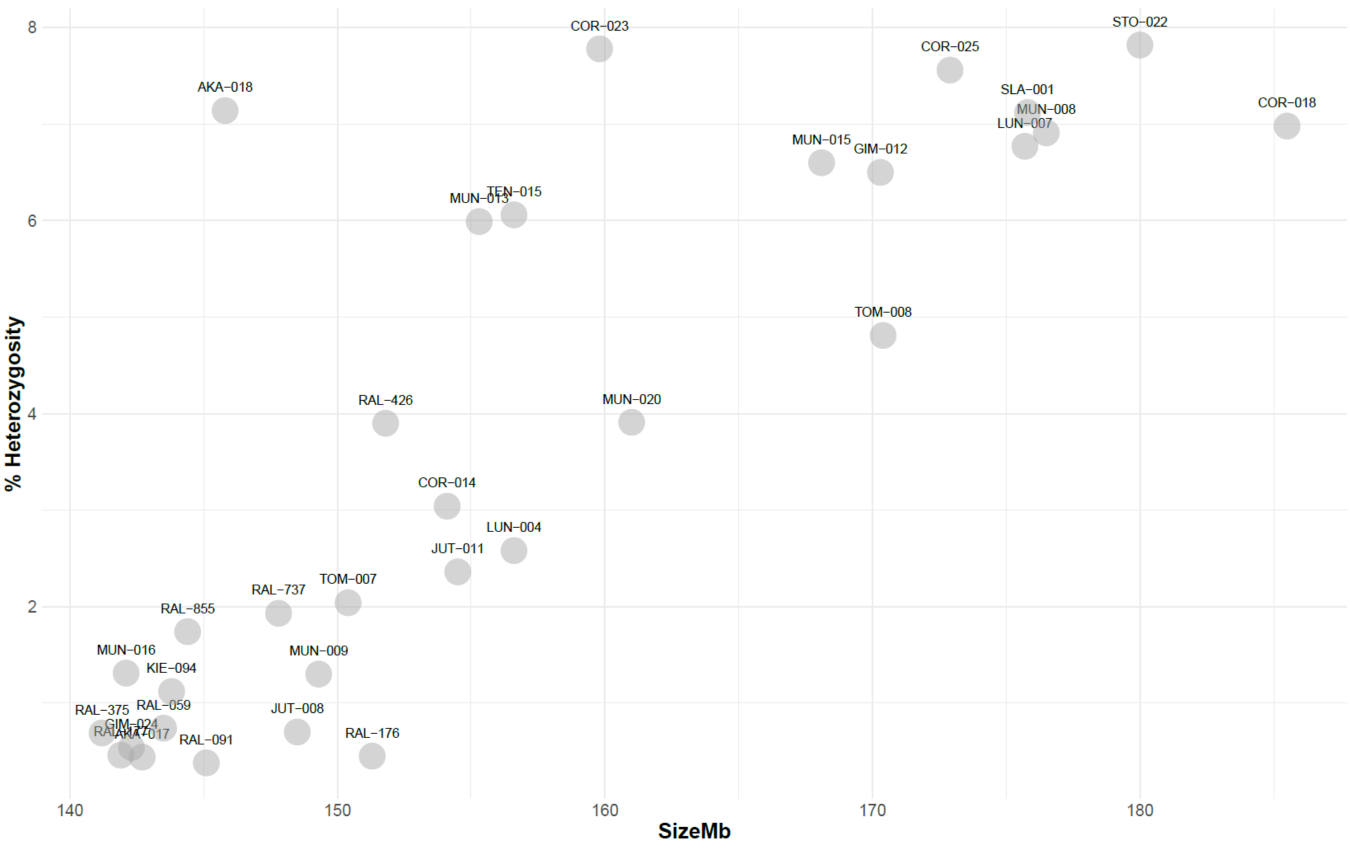

**Figure S17.** Scatter plot showing the relationship between the percentage of *BUSCO* duplicates and the size of the assemblies in Mb (SizeHaplot) after running *purge_haplotigs*.

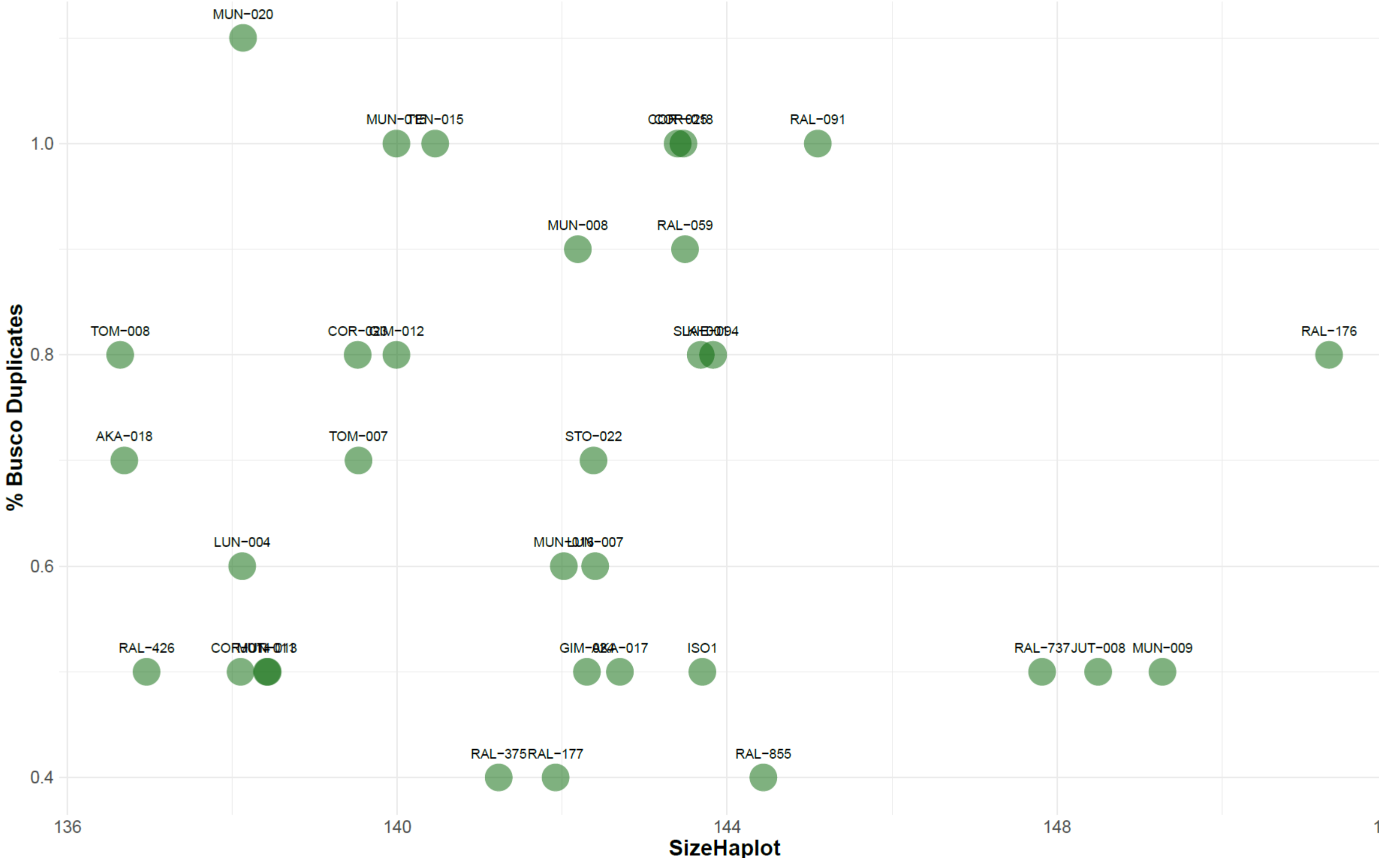

**Figure S18.** Possible outcomes, based on CIGAR information (white box), of the alignment of flanking regions (red blocks, 500 bp) of TE insertions (gray block, 1000 bp in this example) using minimap2. **A)** Non-reference insertion. The flanking regions of the TE align contiguously in the reference genome, which indicates that the TE is absent. **B)** Reference insertion. The CIGAR information shows two alignment regions (500M) separated by a gap of the size of the insertion (one kilobase in this example, intron-like region =1000N). This result indicates that the flanking regions aligned separated by the expected distance given the size of the TE. **C)** Reference insertion with two possible locations. The CIGAR information shows two gaps in the alignment. One of the expected sizes given the presence of TE and another within one of the flanking regions. The algorithm is able to define the correct TE insertion site based on the expected size of the flanking regions.

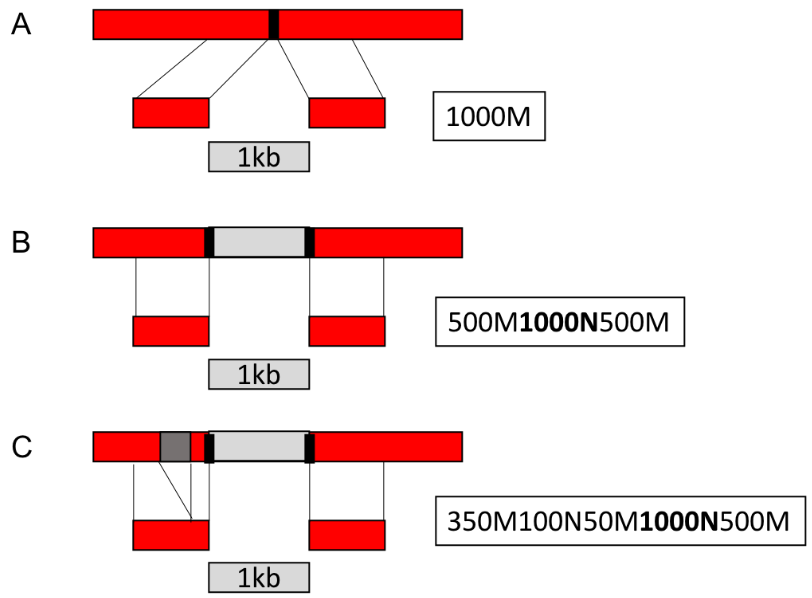

**Figure S19.** TEs in a tandem (gray blocks) transferred by association to reliable upstream and downstream TEs (dark grey blocks, TU: upstream TE; TD: downstream TE). Two conditions must be met to consider that the transfer of a TE is reliable: i) maintain the upstream and downstream genes (blue) in the transference to the reference genome; ii) at least one of their flanking regions (in red) have been uniquely mapped.

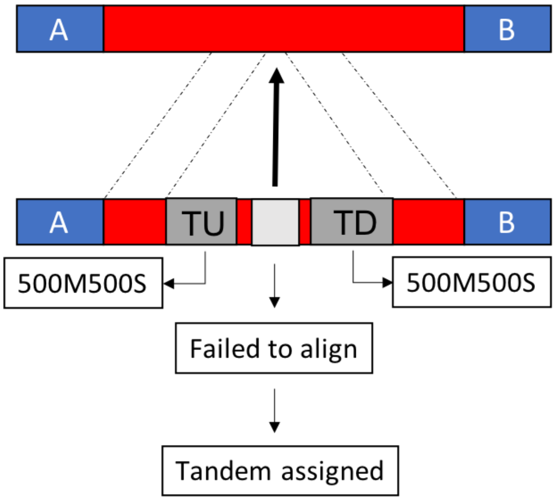

**Figure S20.** TE transference (grey block) to the reference genome is done independently in each strain (G1, G2, G3…G46) by aligning the corresponding TE flanking regions (red blocks) to the reference genome. A) TEs whose transfer coordinates overlapped with a TE present in the reference genome which belongs to the same family are classified as orthologous TEs and are assigned the same name (ID) and coordinates as the TE in the reference genome. B) TEs in different strains that do not have orthologs in the reference genome but that they belong to the same family and their insertion points are less than 50bp from each other are classified as orthologous TEs. The most 5' (P1) and 3' (P2) extreme insertion points are set as the new coordinates and a new identifier (ID) was assigned consisting of the chromosome arm, coordinates and family to which they belong.

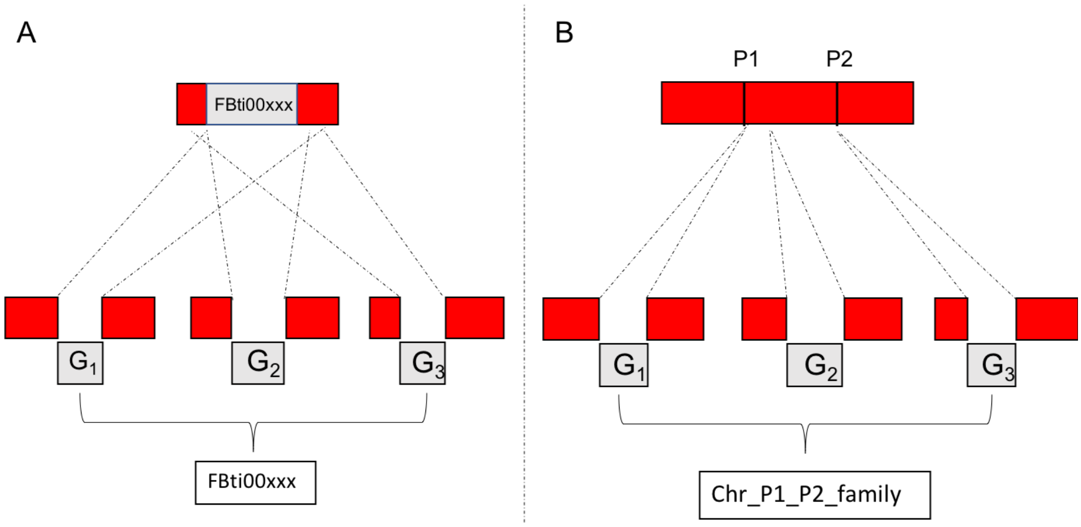

**Figure S21.** PCA plot performed using the PCA module of *QTLTools* on the normalized gene expression matrix (phenotype table). A clear separation between RAL and no-RAL strains is observed.

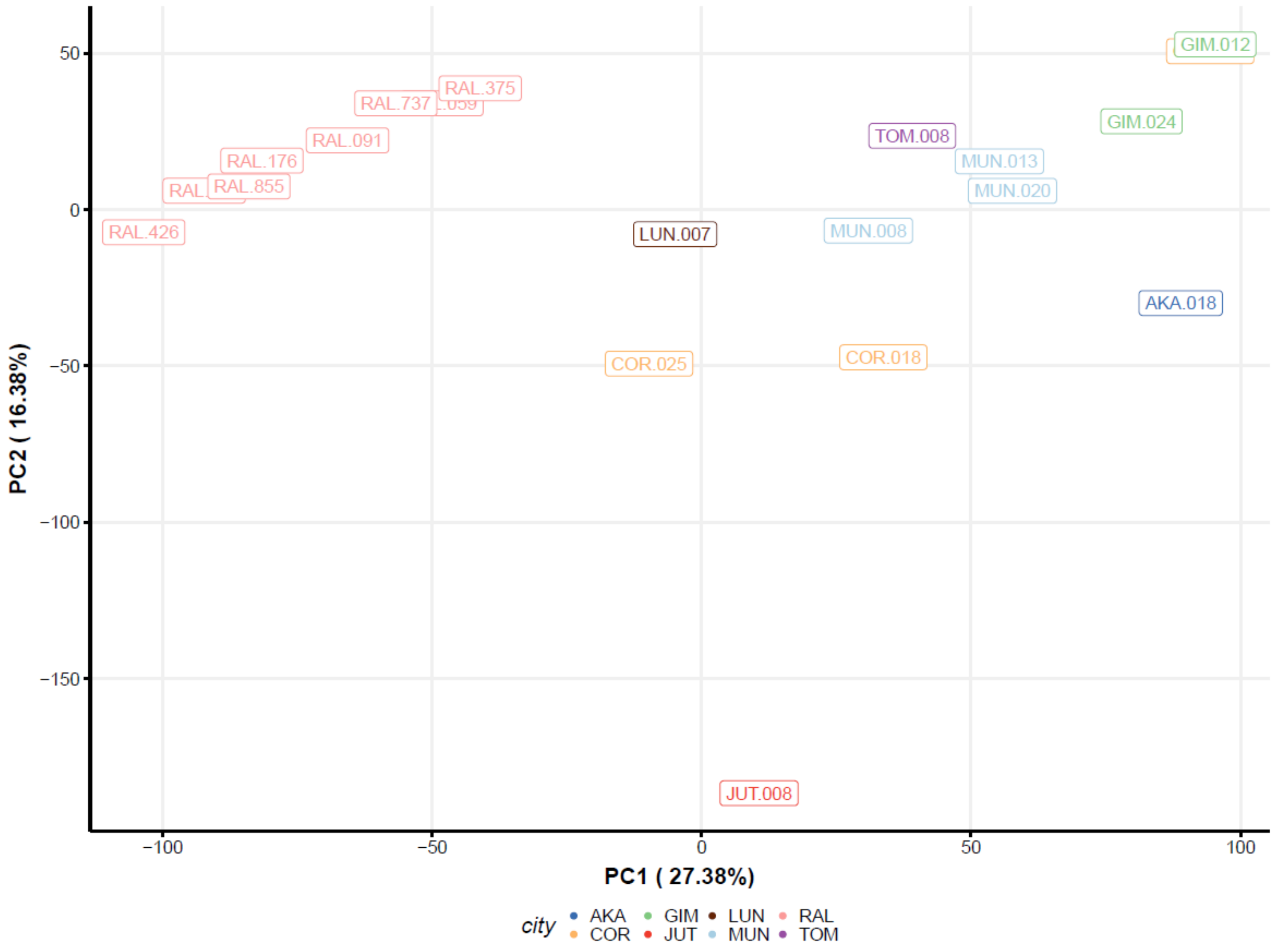

**Figure S22.** Percentage of variation explained for each principal component in the PCA analysis obtained from expression data (Figure S21). The three main principal components (explaining 54.5% of the variation) are added as covariates to the *eQTLTool* model.

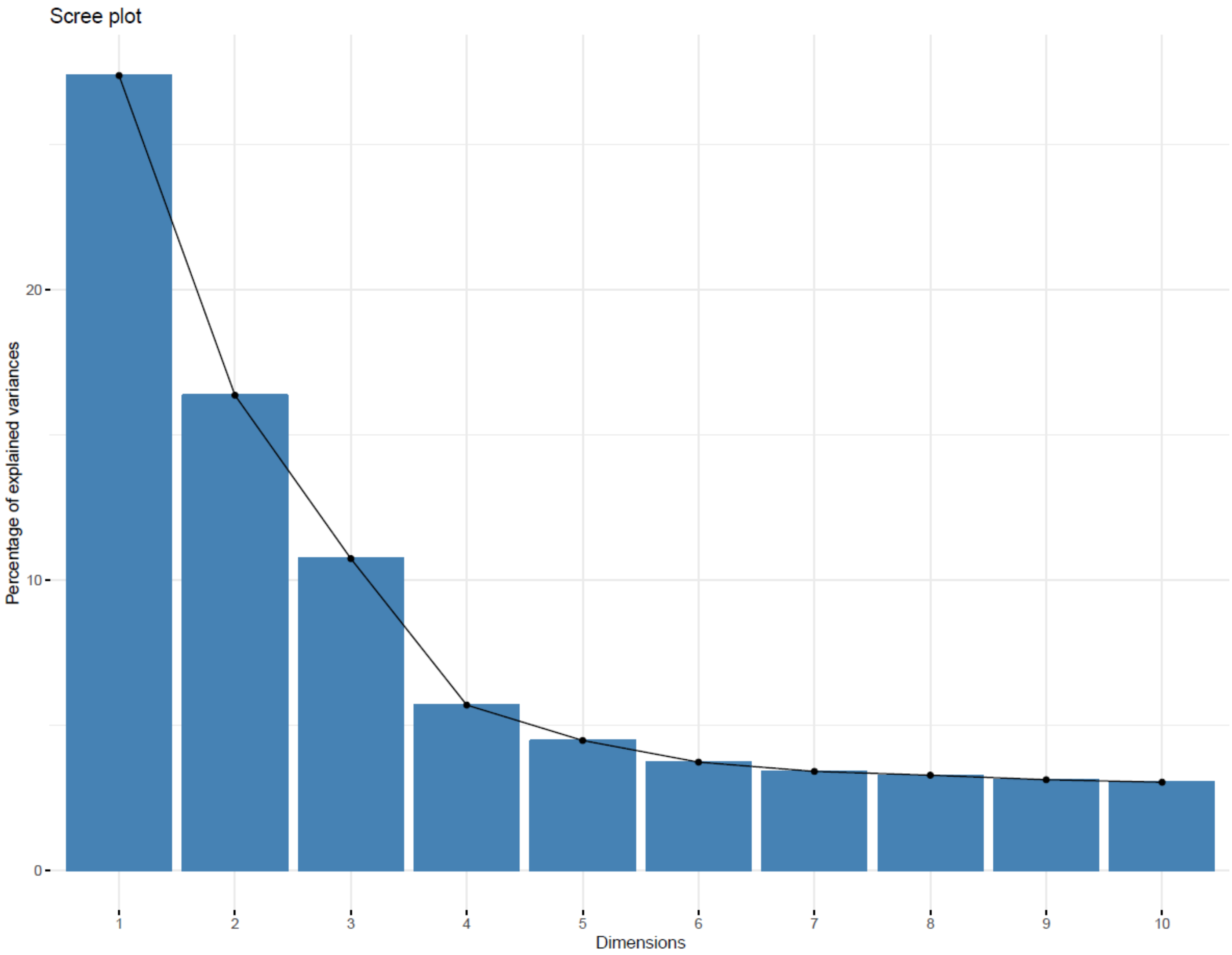

**Figure S23.** Graphical representation of the pairwise matrix of the number of piRNA clusters shared between genomes. Figure and analysis were performed using *intervene* (Khan & Mathelier 2017). Legend at the right indicates de number of shared piRNA clusters.

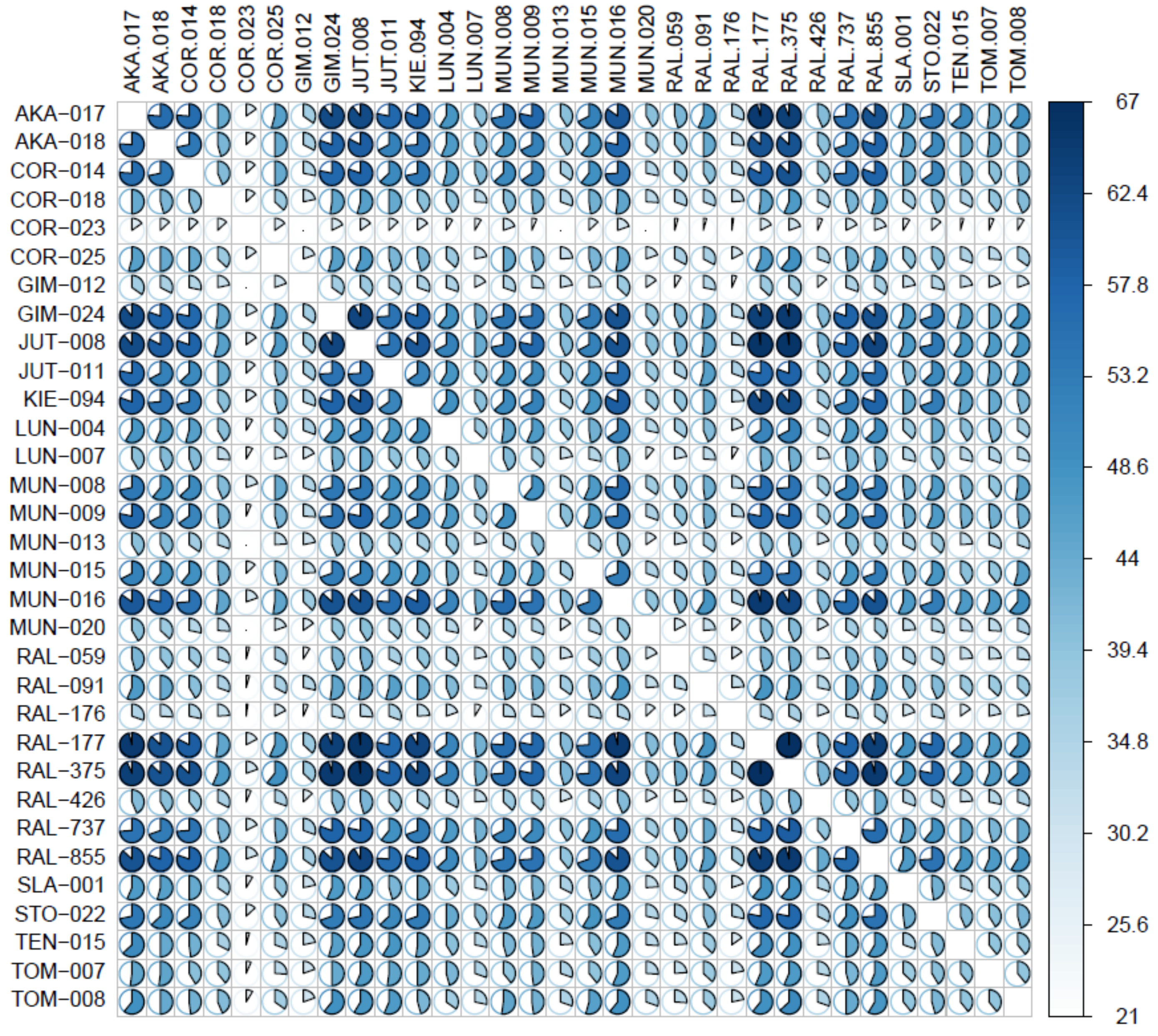
