## Supplementary File S1 for "Population-scale long-read sequencing uncovers transposable elements contributing to gene expression variation and associated with adaptive signatures in *Drosophila melanogaster*"

### Supplementary File 1

#### 1.1 DNA extraction and long-read sequencing

DNA for sequencing using the Oxford Nanopore Technologies (ONT) was extracted from 100 female flies from each strain (except for strain LUN-004, where DNA was extracted from 200 female flies). Two different extraction kits were compared: Gentra Puregene Tissue Kit (Qiagen) followed by a phenol-chlorophorm extraction and the Blood and Cell Culture DNA Mini Kit (Qiagen). The DNA extracted with the Blood and Cell Culture DNA Mini Kit showed better A260/280 and A260/230 ratios and better sequencing throughput, so most of the DNA for Nanopore sequencing was extracted using this kit (Table S2A-B). For the Blood and Cell Culture DNA Mini Kit, we followed manufacturer's instructions but DNA was eluted in half of the recommended buffer volume in order to increase DNA concentration. Briefly, 100 flies from each strain were mechanically homogenized in buffer G2 (RNase and Proteinase K added) and lysed for 2h at 50ºC and DNA was purified in a Qiagen Genomic-tip (20/G), washed and eluted in buffer QF. The DNA was precipitated with isopropanol and resuspended in 15ul of TE buffer. DNA quality of the samples was assessed by measuring DNA concentration using a Qubit® fluorometer and A260/280 and A260/230 ratios in a Nanodrop® spectrophotometer. DNA integrity was assessed in a 1% agarose gel electrophoresis.

Nanopore libraries were constructed using the Ligation Sequencing Kit (SQK-LSK108 or SQK-LSK109, see Table S2A) following manufacturer's instructions. 1-2ug of DNA from each sample was used to start with the library workflow. Sample fragmentation in g-Tubes (Covaris) was evaluated in order to increase sequencing throughput (see fragmentation conditions in Table S2A) but no higher throughput was observed. Thus, most of the samples were sequenced without DNA fragmentation. DNA concentration was assessed during all the procedure to ensure we had enough DNA for sequencing. The libraries were run using the MinION device with R9.4 flow cells for 48 hours at -180mV. The total number of flow cells used for each strain is given in Table S2C. If more than one sample was run in a single flowcell, we used the Native Barcoding Expansion 1-12 together with the Ligation Sequencing Kit, and equal amounts of each sample were loaded in the flow cell.

#### 1.2 Assembly of genomes sequenced with ONT

Genome assemblies for ONT sequenced genomes started by applying *Canu* (v.1.7) (Koren *et al.* 2017) over the raw fastq sequences with default options. We then used *minimap2* (v.2.9) (Li 2018) for mapping fastq files back to the draft assembly and then we used *Racon* (v.1.0) (Vaser *et al.* 2017) to generate the genomic consensus. We repeated the *minimap2+Racon* process four times, since we observed that a fifth round was not significantly improving the assembly resolution, measured as the percentage of complete *BUSCO* genes (see below). Then, we mapped raw reads over the Racon-polished genomes and used *Nanopolish* (v.0.10.1) (https://github.com/jts/nanopolish) to call an improved consensus sequence (Figure S12). Since the quality of the polished assemblies was still below PacBio assemblies, we decided to use Illumina short-read sequences for a final polishing step. In order to do that, we first removed adapters and low quality sequences from Illumina reads using *cutadapt* (v1.16) (Martin 2011) and mapped the trimmed reads to the *Racon+Nanopolish* assembly using *BWA* (v.0.7.12) (http://bio-bwa.sourceforge.net/). Resulting BAM files were imported in *Pilon* (v.1.22) (Walker *et al.* 2014) for obtaining final polished assemblies (Figure S12), which greatly enhance the resolution of the genomes, as shown by the increased percentage of complete *BUSCO* genes, making them comparable in quality to the ones obtained with PacBio sequencing (Table 1, Table S2C).

#### 1.3 Genome deduplication

We found that the percentage of repeats estimated by *RepeatMasker* in the genomes positively correlated with the assembly size (r=0.86, p-value=8.8e-11, Figure S13) and negatively correlated with genome contiguity (N50) (r=-0.50, p-value=0.002, Figure S14). To further investigate possible causes of the variation in genome size we checked the percentage of duplicated regions in the genome using the duplicated *BUSCO* genes as a proxy. We found percentages of duplication between 0.5% and 7.8% and a strong positive correlation between the genome size and the percentage of *BUSCO* duplicates (r=0.95, p-value=2.2e-16, Figure S15).

Genomic regional duplications could be explained by the presence of heterozygous regions in the genome that could not be consolidated in a single allelic variant during the assembly (Roach *et al.* 2018). We calculated whole genome heterozygosity for the 32 genomes under study using short-read sequences. For ONT genomes, we mapped the Illumina short-reads to the ISO1 genome using the *GATK* (v4.0) (McKenna *et al.* 2010) best practices for variant discovery (Van der Auwera *et al.* 2013). Then, we used the *bcftools stats* (v1.9) (Li 2011) for calculating the percentage of heterozygous SNPs. For PacBio genomes we generated Illumina-like reads from the corrected PacBio reads using *randomreads.sh* from *BBTools* (v.38.23) (Bushnell 2014). We found a strong correlation between the estimated heterozygosity and the assembly size (r=0.82, p-value=5.9e-09, Figure S16).

Since the level of heterozygosity is one of the main determinants for assembly quality, we used *purge_haplotigs* (Roach *et al.* 2018) for identifying pairs of contigs that were syntenic to eventually keep only one in the (haploid) assembly. We first evaluated the effects of *purge_haplotigs* (Roach *et al.* 2018) over the genomes with different levels of heterozygosity and we found that for levels lower than 0.2% the duplicated *BUSCO* scores were not decreasing, so we decided to run it only for genomes showing levels of heterozygosity > 0.2% (JUT-011, LUN-004, COR-014, TEN-015, SLA-001, STO-022, TOM-007, MUN-008, COR-025, MUN-013, COR-023, LUN-007, GIM-012, TOM-008, COR-018, MUN-015, RAL426, AKA-018, MUN-020). When considering the haploid assembly, the duplicated *BUSCO* genes significantly decreased to 0.4%-1.1% (Table 1) and the correlation with the genome size disappeared (r=0.06, p-value=0.73, Figure S17), while the complete *BUSCO* genes remained around 98% on average, showing that the vast majority of the genome was represented in the *de novo* haploid assemblies (Table 1).

#### 1.4 Gene transfer

Gene transfer, *i.e.* determining gene coordinates in the *de novo* assemblies, was performed by mapping the coding sequence (CDS) of the 13,836 protein-coding genes annotated in the *Drosophila melanogaster* reference genome (v. 6.31) against each of the sequenced genomes. Briefly, for each protein-coding gene, the CDS of the longest transcript was obtained using the *BioPython* module (Cock *et al.* 2009) and then aligned to the genomes of each strain using *minimap2* (v.2.9) (Li 2018) with the *splice* mode (*minimap2* rates an alignment by the score of the max-scoring sub-segment, *excluding* introns). After that, we defined the mapping coordinates of each gene, using the *pysam* library (https://github.com/pysam-developers/pysam) (Li *et al.* 2009). In cases where the CDS did not map or map in multiple locations in the *de novo* genomes, the complete gene sequence was used to define the coordinates of the gene in the genome of interest. In order to do this, we used *minimap2* but using the *asm5* mode (no splicing allowed), and the exact mapping coordinates of the CDS were defined through the implementation of the hamming distance algorithm.

#### 1.5 Calculation of piRNA Clusters BUSCO (CUSCO) score

*CUSCO* quality metric determines the percentage of contiguously assembled piRNA clusters in a genome assembly. This metric, unlike most traditional genome assembly metrics such as gene BUSCO or the N50, provides information about the genome quality in the context of TEs (Wierzbicki *et al.* 2020). We used the scripts and the piRNA flanking regions (Brennecke *et al.* 2007) available at https://sourceforge.net/projects/cuscoquality/ for calculating CUSCO values in the 32 sequenced genomes (Table S3A). Two different BUSCO scores were calculated, the c.CUSCO that determines the percentage of contiguously assembled piRNA clusters at the contig level and the sc.CUSCO that determines such percentage at the scaffold level (e.g stretches of 'Ns' are allowed). Overall, 80 (94%) out of the 85 piRNA clusters analyzed were detected in at least one genome assembly and seven clusters were identified in the 32 genomes (Table S3B). As expected, the number of genomes in which a piRNA cluster is detected is negatively correlated with the size of the cluster (r= -0.47), suggesting that as the larger the cluster, the more difficult to get it contiguously assembled (Figure S2). Average pairwise number of shared piRNAs between genomes was 42.5, with genomes RAL-375 and RAL-177 showing the highest number of shared piRNA clusters (67) and COR-023 vs GIM-012, MUN-013 and MUN-020 with the lowest number of common clusters (21) (Figure S23).

In an attempt to determine methodological or biological causes explaining the variability in the CUSCO values, we tested different linear models considering several putative explanatory variables. On one side, we analyzed three pre-assembly metrics: heterozygosity, sequencing coverage and the N50 of the sequenced reads. On the other side, we considered five assembly metrics: the contigs N50, number of contigs in the assembly, size of the final genome assembly and the BUSCO complete and duplicated scores. We found that among the pre-assembly metrics, the N50 of the reads and the levels of heterozygosity seem to be the main factors explaining CUSCO scores (Table S3C). Moreover, regarding the assembly metrics, while the N50 of the contigs is significant, the total number of contigs and the size of the assembly seems to explain much better the observed CUSCO values (Table S3C). Interestingly, and in agreement with the previously described (Wierzbicki *et al.* 2020), neither the complete not the duplicated BUSCO scores are good predictors of the CUSCO score (Table S3C).

#### 1.6 Construction of the Manually Curated TE (MCTE) library

The creation and curation of the MCTE library involved four main steps: i) running the *TEdenovo* pipeline (default parameters) over 13 out of the 32 pre-scaffolded genomes for discovering new consensus TE sequences; (ii) clustering and redundancy removal; (iii) removing artifactual sequences; and (iv) classification of consensus sequences into TE families. Note that steps 2-4 were performed in an interweaved and no consecutive way. Briefly, once we obtained the consensus sequences with *TEdenovo,* as a first filtering step we run *TEannot* using the whole *de novo* library for each genome over the euchromatic region of the corresponding genome and we kept only the Full Length Fragment (FLF) consensus as classified by *REPET* (i.e. fragment covering more than 95% of the consensus sequence). From the 2,319 FLF sequences, we discarded 495 by visual exploration of the copies annotated with each consensus using the *plotCoverage* tool from *REPET*, discarding those consensuses showing a high number of small copies. For the remaining 1,824 FLF sequences we attempted to assign them to known TE families by aligning them against the library of 179 curated canonical sequences of Drosophila TEs from the Berkeley Drosophila Genome Project (BDGP, v.9.4.1, https://fruitfly.org/p_disrupt/TE.html), 127 of which correspond to *D. melanogaster* TE families and the remaining 52 to TE families from other Drosophila species. We used *BLAT* (v.35) (Kent 2002) with default parameters for assigning FLFs to families in the BDGP set. We considered a consensus sequence to belong to a BDGP family when they were at least 90% identical. If there was tie in the identity values, we choose the sequence with the higher breadth of coverage (covered length) when considering all the aligned sections. We found 1,374 FLFs consensuses with significant matches against 117 BDGP sequences. For the FLFs matching the 117 BDGP we performed Multiple Sequence Alignments (MSA) of the FLF from all genomes plus the BDGP, sequence in order to obtain a consensus sequence representative of all genomes. These 117 consensus sequences were added to our MCTE library as representative sequences of the TE family to which they matched in the BDGP (Table S5). Moreover, there were 14 *D. melanogaster* BDGP sequences without any match against the list of FLF, however we also added these 14 sequences to the MCTE library.

Moreover, in order to look for families not represented in the BDGP library, we further explored two sets of *de novo* consensus sequences: the 450 FLFs that were not matching with any BDGP sequences and the 25,690 non-FLFs. In the latter case, we first visually explored the copies annotated with each non-FLFs (using the *plotCoverage* tool in *REPET*), and we kept only those showing three or more long copies and not showing predominantly short copies, which resulted in only 337 candidates (good non-FLF sequences). Then, we discarded those sequences with high similarity to sequences already present in the MCTE library by excluding those with identities > 90% according to *BLAT* (v.35), resulting in 98 non-FLF putative new sequences that were clustered together with the unknown 450 unclassified FLF using *MCL* (Dongen 2000) through the *LaunchMCL.py* script from *REPET*. From the 548 sequences, we obtained 58 clusters plus 60 sequences that were not clustered with any other sequence. For the 58 clusters, we performed MSAs and generated the consensus sequences. These 58 consensus sequences plus the 60 unclustered were further analyzed to identify artifacts, families not present in the BDGP or new families not present in any database. Artifactual consensus were identified by manual inspection of the *PASTEC* (v2.0) (Hoede *et al.* 2014) annotations over each sequence. We discarded those sequences that either were chimeric combinations of more than one TE family, were containing an ORF from a non-TE associated transcript, or those for which we observed multiple simple sequence repeats (SSR) annotations all along the sequence. After manual inspection, 16 sequences remained from the 58 clusters (noBDGPmclCluster) and 18 from the unclustered sequences (noBDGPunClustered) (Table S5). To further characterize these 34 consensuses, we used *RepeatMasker* (v.4) (Smit 2015) with the release *RepBaseRepeatMaskerEdition-20181026* of *RepBase* (Bao et al. 2015) and the parameter *-species insects* in order to assign families to these members of the MCTE library. 24 of the unclassified consensuses showed similarities with sequences in *RepBase*. For the remaining 10 sequences we performed *tblastx* searches against the consensus sequences that were previously assigned to a family using the BDGP strategy. Seven sequences showed significant similarities with some of these consensus sequences. Overall, 31 of the 34 unclassified consensuses were assigned to TE families by either the *RepBase* or the *tblastx* strategy, including seven with similarities with TE families from *D. simulans*, eight with TE families from other Drosophila species, one with a TE family from *C. amoena* and 15 with TE families from *D. melanogaster*. The remaining three consensus sequences showing no similarities either with BDGP nor *RepBase*, were further evaluated for its potential as new unknown families.

##### 1.7 MCTE library performance

In order to determine the performance of the *TEannot* pipeline using the MCTE library we compared our TE annotation strategy with the current annotation of 1,301 euchromatic TEs available in FlyBase (Gramates *et al.* 2017). We first compared the number of copies per family from each annotation. 95 out of the 146 families represented in the MCTE library were present in both annotations (Table S6A, Figure 2A). Only two families (*frogger* and *gypsy3*), with one copy each in FlyBase, were not identified using *REPET*. In addition, four copies annotated in FlyBase as members of the *S2* family were annotated in *REPET* as *S-element*, since the *S2* consensus was removed from our MCTE library for being redundant with the *S-element* consensus. For 28 families, we found at least one copy using *REPET* annotation while no copies were found in the FlyBase annotation. Some of them correspond to families not present in the whole FlyBase annotation (e.g. *LARD*, *Kepler* and *THARE*) but others are families that are only observed in the heterochromatic regions according to the FlyBase annotation (e.g. *gypsy10*, *BS4* and *ZAM*) (Table S6A, Figure 2A). For those families identified by both annotations, we observed no significant differences in the number of copies among *REPET* and FlyBase annotations (FDR p-value > 0.05, $X^2$ test, Table S6A, Figure 2A), with exception of the *INE-1* elements, for which *REPET* annotated a proportionally larger number of copies than FlyBase (adjusted p-value < 0.0001, $X^2$ test, Table S6A).

At the coordinates level, we compared the overlap between coordinates predicted by *REPET* and the ones annotated in FlyBase using *BEDtools intersect* (v2.25.0) when considering different percentages of overlap (reciprocal minimum breadth of coverage, Table S6B). Note that for this analysis, we excluded TEs shorter than 100bp, TEs from the *INE-1* family and nested TEs. TEs from the *INE-1* family were excluded because they are usually short, highly variable, are not flanked by target site duplications and their termini lack direct or inverted repeats (Locke *et al.* 1999), which makes extremely difficult to determine the correct start and end coordinates. We end up with 884 TEs from FlyBase and 1,719 TEs from the *REPET* annotation, for which we found that 84.62% of the FlyBase annotations were overlapping at least at the 95% of the copy length with *REPET* annotations (Figure 2B, Table S6B). Moreover, overall sensitivity and specificity of *REPET* annotation when comparing with FlyBase were 99.44 and 99.29, respectively, as calculated according to (Quesneville *et al.* 2005) (Table S6C).

We also observed that the average copy length was significantly smaller in *REPET* annotations than in FlyBase (1,952bp and 2,769bp, respectively, p-value = 9.7E-12, t-test). This was also reflected in the differences of the distributions of the length of the copies (p-value = 1.9E-12, Kolmogorov-Smirnov's test, Figure 2C), with a clear enrichment of small copies in our *REPET* annotation. In fact, if we consider only copies > 2Kb in both annotations, the distribution of sizes is not statistically different (p-value = 0.1757, Kolmogorov-Smirnov's test, Figure 2C).

Finally, in order to determine the performance of the MCTE library we compared the annotation obtained with the MCTE library and the current *D. melanogaster* consensus sequences present in the BDGP using *RepeatMasker* (v.4) (Smit 2015) for 13 genomes utilized for building the MCTE plus the ISO-1 genome . After running *RepeatMasker* with each library over the 14 genomes, we used *BEDtools coverage* (v2.25.0) for determining the breadth of coverage of annotation using the MCTE on annotation using the BDGP.

#### 1.8 Characterization of unknown TE families in other Drosophila species

After construction of the MCTE library (see above), three consensus sequences showing no similarities with any other sequence in the BDGP library or in *RepBase* were further analyzed to determine whether they could represent unknown TE families in *D. melanogaster*. In order to determine whether these elements where present in other Drosophila species, we performed *de novo* genome assembly and *de novo* TE annotation of five out of the 15 Drosophila genomes recently sequenced using ONT (Miller *et al.* 2018). We selected five species based on their widest phylogenetic distribution (*D. simulans, D. yakuba, D. virilis, D. bipectinata* and *D. pseudooscura*). Genome assembly, polishing and *de novo* TE prediction were performed following the same pipeline described for *D. melanogaster* genomes (see above). *De novo* TE annotation of the five genomes generated a library of 9,574 consensus sequences that were used as database for searching similarities with the three new family consensuses through *tblastx* searches.

#### 1.9 Identification of inverted repeats

We extracted the coordinates of all copies annotated with the newly identified *TIR* consensus sequence in all genomes. Then, we extend such coordinates 1Kb using *BEDtools slop* (v2.25.0) and extracted the corresponding fasta sequences using *BEDtools getfasta* (v2.25.0). For identifying inverted repeats in such sequences we used *EMBOSS palindrome* (v6.6.0) (Rice *et al.* 2000) with the following parameters: -minpallen 50 -maxpallen 1000 -gaplimit 10000 -nummismatches 20. Results were then manually explored to determine whether the palindrome detected regions were at the ends of the provided fasta sequences.

#### 1.10 Comparison with short-read TE annotation methods

We used two different software that use short-read sequencing to detect non-reference TE insertions (present in at least one sample and absence in the reference genome): *TEMP* (v.1.05) (Zhuang et al. 2014) and *TIDAL* (v.1.0) (Rahman *et al.* 2015). These two software were used to detect euchromatic non-reference TE insertions in 11 strains representative of the geographic variability of our samples that are among the most complete genomes (Table 1). In both cases, we also used the MCTE library as input source for TE annotation. *TEMP* software uses the information provided by both pair-end and split reads to infer the TE insertion interval at nucleotide resolution, while *TIDAL* was designed to be run using single-end and split reads (the *TIDAL* pipeline automatically converts all pair-end into single-end reads). Before running *TEMP*, we mapped the pair-end sequencing reads using the *bwa mem* algorithm of *BWA* (v.0.7.16a) to the reference *D. melanogaster* genome (v.6). After mapping, we filtered the data to remove duplicates and realigned around indels using *picard* (v.2.8.3), *GATK* (v.3 3.7) and *SAMtools* (v.1.6) for indexing. The *TEMP* module insertion was run with command line options -m 3 -c 8 -x 30 and -f 500 and -f 300 for European and North American strains, respectively. *TIDAL* was run using default settings (Rahman *et al.* 2015). For 10 out of the 11 strains, *TIDAL* was run on *fastq* files using *TIDAL_from_fastq.sh* script and only for RAL-375, *TIDAL* was run on SRA files using *TIDAL_from_sra_DGRP.sh* script since that particular strain has multiple libraries with varying length concatenated into one SRA entry. The *min_len* parameter was set to 100 for all the strains except for RAL-177 and RAL-375 in which this parameter was set to 95 and 75, respectively.

For *TEMP* results and to avoid including in the analysis predictions of TE insertion that might be false positives, we only considered as reliable those TE insertions with split-read support at both ends of the insertion ("1p1"). All predictions made by *TIDAL* were considered as reliable since it uses a *BLAT* score for filtering the potentially false positives. We filtered out those TE insertions (estimated by *REPET*, *TIDAL* and *TEMP*) that were in the heterochromatic regions.

Next, we concatenated the TEs found in each strain in a single file while adding a specific ID corresponding to the name of the strain where the TE insertion was predicted. This was done for each method (*REPET*, *TIDAL* and *TEMP*) separately and adding a second ID corresponding to the method with which the TE insertion was predicted. The three concatenated files were in turn concatenated in a single file. *BEDtools* (v.2.18) (Quinlan & Hall 2010) was used with options *sortBed* and *merge* to sort the TEs and to merge overlapping TE intervals between the 11 strains found with the three different methods while keeping information about the strains and the method with which the TE insertion was predicted. *R* (v.3.5.1) was used to compute the number of TEs found by only one, two, or three methods (Table S8A). There were some few cases in which a particular TE found by one method overlaps with more than one TE found by another method. Those TEs were removed from the final results.

#### 1.11 TE orthology identification

To identify orthologous TEs among the sequenced genomes we transferred the TE coordinates from each strain to the ISO1 reference genome. In order to do that, we used *minimap2* (v.2.9) (Li 2018) to map both the TE sequence and their 500 flanking nucleotides to the ISO1 reference genome using the *asm5* and the *splice* mode, respectively. TEs having stretches of N's (scaffolds joining) at both flanking regions were not transferred. Briefly, we first mapped TE flanking regions from each genome using *minimap2* in *splice* mode and classified the alignments as unique, multi-mapped or not mapped. In case of unique mappings, we observed three different cases. First, TEs whose flanking regions aligned completely or almost completely to the reference genome (the 500bp flanking sequence on both sides mapped together in the genome) were defined as *de novo* TE insertions regarding the reference genome (they are not present in the reference genome). Their exact breakpoint (insertion site) was defined by the information present at the CIGAR string of the *minimap2* alignment result (i.e. the number of insertions, deletions, etc., Figure S18A). Second, TEs whose flanking regions aligned separately in the genome (having an intron-like insertion between them; represented in the CIGAR string as Ns). For those, the reference genome coordinates were defined by the start and end coordinates of the intron-like region. These TEs were assumed to be present in the reference genome (Figure S18B). Third, TEs whose flanking region align with more than one intronic-like region. In this case, we kept the intron-like insertion that separates both flanking regions (for example, keeping the intron-like insertion flanked by 500bp at one of its ends). These TEs were also assumed to be present in the reference genome (Figure S18C).

TEs that could not be unambiguously defined based on the strategy described above (i.e. more than two intron-like regions, soft-clipping bases, inconsistent TE size regarding the size expected from the alignment) were subjected to a second round of assessment using the multi-mapping flanking regions. To define their coordinates in the reference genome, we used the information of the alignment of the complete sequence of the TE (*minimap2*, *asm5* mode). If the TE sequence mapped to a unique location, we defined as reference coordinates those that overlapped with the coordinates defined by the flanking regions. When the complete sequence of the TE aligned to multiple locations, the reference coordinates were defined based on the location showing less mismatches. Finally, when the TE did not align at all, we defined its position based on the distance to the closest transferred gene. We tagged these TEs as ambiguously transferred TEs.

TEs were also classified as nested or tandem if were completely located within another TE or if a TE was located to less than 500bp either upstream or downstream of another TE. Nested or tandem TEs are more difficult to transfer because part of their flanking regions are repetitive regions that failed to align in the reference genome or are misaligned. Therefore, in order to properly transfer these TEs, we rely on the location of their associated TE, whether it is in tandem or nested, where there is evidence that their transfer was correct. Thus, we define a set of reliable TEs, which must be uniquely mapped and maintain the gene synteny in the transfer (i.e., genes at the flanking regions of the TE are the same in both, the reference and the genome of interest). We used *BEDtools closest* (v2.25.0) to check whether the synteny was conserved by looking the closest upstream and downstream genes. So, in a case where a TE could not be correctly transferred and it was classified as nested or in tandem, we defined its insertion coordinates as the position one base next to the associated TE from the reliable set (Figure S19). Finally, we discarded from the transference those TEs where the coordinates were not clearly defined and their location could not be redefined based on a reliable TE.

Once the TEs of each strain were transferred to the reference genome, we used these coordinates to find orthologous TEs between the different strains using *BEDtools intersect* (v2.25.0). If a TE of a given strain was present in the reference genome ISO1 and belonged to the same Family, we kept the TE ID and coordinates of the TE in the reference genome (Figure S20A). If that TE was not found in the ISO1, the reference coordinates were re-established as a single insertion point and classified as a non-reference TEs. In addition, to establish the new coordinates and ID of each TE the following considerations were taken: i) if a TE intersects with both a FlyBase and a *REPET* annotation, only FlyBase coordinates were assigned; ii) if a TE intersects with more than one FlyBase TE, a new set of coordinates matching the ones in FlyBase was created for each intersection; iii) if a TE only intersects with the *REPET* annotation, *REPET* coordinates and ID were assigned to that TE. On the other hand, we considered that non-reference TEs were orthologous if they belong to the same TE family and the distance between the predicted insertion points in the different strains were less than 50bp (transitive associations). We used *BEDtools merge* (v2.25.0) in order to find non-reference orthologous TEs and their assigned insertion site were determined by the most 5' (P1) and 3' (P2) extreme positions among the predicted insertion points of the orthologous TE in each strain. We assigned a new ID consisting of the chromosome arm, coordinates (P1 and P2) and the family to which they belong (Figure S20B).

##### 1.12 Selection Analysis

We evaluated whether TEs showed evidences of positive selection using the orthologous information for the 32 genomes sequenced in this work plus 14 genomes previously sequenced (Chakraborty *et al.* 2019) and Single Nucleotide Polymorphisms (SNPs) as a proxy analyzed with *selscan* (v1.2.0a) (Szpiech & Hernandez 2014). Briefly, *selscan* uses as input a variant file containing genetic variant information and a *map* file containing information about the genetic and physical positions for each variant. SNPs were called using the *GATK* (v4.0) (McKenna *et al.* 2010) *HaplotypeCaller* best practices for variant discovery (Van der Auwera *et al.* 2013) over the alignments generated by mapping to the ISO1 either, the Illumina short-reads (for genomes sequenced by ONT) or Illumina-like reads generated using *randomreads.sh* from *BBTools* (Bushnell 2014) from the corrected PacBio reads. After running the *GATK HaplotypeCaller* for each genome, we merged them using the *CombineGVCFs* command and we performed the joint genotyping using *GenotypeGVCFs*. We kept only biallelic SNPs using the GATK command *SelectVariants* (parameters *-select-type SNP --restrict-alleles-to BIALLELIC*). Finally, we removed SNPs with *missing data* in at least one genome, resulting in a total of 2,797,589 SNPs (available at http://dx.doi.org/10.20350/digitalCSIC/13708). Since *selscan* methods assume phased haplotypes, we used *SHAPEIT4* (v.4.1) (Delaneau *et al.* 2019) for determining haplotypes in the SNP data. We adapted the *vcf* format to that expected by *SHAPEIT4* and we created a genetic map file based on the recombination rates calculated by (Comeron *et al.* 2012) and the genetic positions available in FlyBase (https://wiki.flybase.org/wiki/FlyBase:Maps, last updated June 15, 2016). We then indexed *vcf* files using *bcftools index* (v1.9) (Li 2011) and run *SHAPEIT4* for each chromosomal arm separately. We then used *selscan* to calculate three statistics designed to identify putative regions under recent or ongoing positive selection for all positions for the three classes of *vcfs*. Specifically, we run iHS (Voight *et al.* 2006), iHH12 (Garud *et al.* 2015; Torres *et al.* 2018) and nSL (Ferrer-Admetlla *et al.* 2014). Unstandardized iHS, iHH12 and nSL values were normalized in 100 frequency bins across the entire chromosome using *norm*, a package provided with *selscan*. We determined the significance of the selection statistics by comparing the normalized values with the empirical distribution of values for SNPs falling within the first 8–30 base pairs of small introns (<=65bp) that are considered to be neutrally evolving (Parsch *et al.* 2010). For each statistic and each chromosomal arm, we determined the critical values as the 5$^{th}$ percentile in the case of iHS and nSL and the 95$^{th}$ percentile in the case of iHH12 of the distribution of normalized values of SNPs falling in such neutrally evolving regions (Table S13B). In order to identify TEs putatively linked to the selective sweeps, we analyzed the co-occurrence (in the same strains) of the allele showing signatures of a selective sweep and a nearby TE (<1Kb). Then, for each SNP-TE pair we stablished criteria of 'co-occurrence' by requesting certain number of the strains containing both the SNP allele undergoing a selective sweep and the nearby TE: for TEs present in ≥7 strains we request ≥50% of strains to contain both the significant SNP and the nearby TE, for TEs present in 5-6 strains we request at least 4 of the strains to contain both the allele undergoing a selective sweep and the nearby TE. In all cases, we also requested the TE to be absent in all strains that do not contain the significant SNP (Table S13A).
